# Critical Evaluation of Sphingolipids Detection by MALDI-MSI

**DOI:** 10.1101/2025.02.04.636486

**Authors:** Bo Chen, Ranjana Pathak, Albar Subekti, Xiaolu Cheng, Sukhjit Singh, Anne G. Ostermeyer-Fay, Yusuf A. Hannun, Chiara Luberto, Daniel Canals

## Abstract

The increasing interest in the role of sphingolipids in (patho)physiology has led to the demand for visualization of these lipids within tissue samples (both from animal models and patient specimens) using techniques such as matrix-assisted laser desorption/ionization mass spectrometry imaging (MALDI-MSI). While increasingly adopted, detection of sphingolipids with MALDI-MSI is challenging due to: i) the significant structural variations of sphingolipid molecules, ii) the potential breakdown of the more complex molecules into structurally simpler species which may confound the analysis, and iii) the great difference in levels among sphingolipid classes and subspecies, with the low-abundant ones often being close to the detection limit. In this study, we adopted a multi-pronged approach to establish a robust pipeline for the detection of sphingolipids by MALDI-MSI and to establish best practices and limitations of this technology. First, we evaluated the more commonly adopted methods [2,5-Dihydroxyacetophenon (DHA) or 2,5-Dihydroxybenzoic acid (DHB) matrix in positive ion mode and 1,5-Diaminonaphthalene (DAN) matrix in negative ion mode] using MALDI-MS on reference standards. These standards were used at ratios similar to their relative levels in biological samples to evaluate signal artifacts originating from fragmentation of more complex sphingolipids and impacting low level species. Next, by applying the most appropriate protocol for each sphingolipid class, MALDI-MSI signals were validated in cell culture by modulating specific sphingolipid species using sphingolipid enzymes and inhibitors. Finally, the optimized parameters were utilized on breast cancer tissue from the PyMT mouse model.

We report the optimal signal for sphingomyelin (SM) and, for the first time, Sph in DHB positive ion mode (in cells and PyMT tissue), and the validated detection of ceramides and glycosphingolipids in DAN negative ion mode. We document the extensive fragmentation of SM into sphingosine-1-phosphate (S1P) and even more so into ceramide-1-phosphate (C1P) using DAN in negative ion mode and its effect in generating an artifactual C1P tissue signal; we also report the lack of detectable signal for S1P and C1P in biological samples (cells and tissue) using the more suitable DHB positive ion mode protocol.

## INTRODUCTION

Sphingolipids are structural components of cellular membranes that play a pivotal role in cellular signaling and metabolism, extensively investigated in the context of cancer progression [1, 2]. Structurally, sphingolipid species are formed by a sphingoid building block, typically **sphingosine (Sph)**. N-acylation of Sph with various CoA-activated fatty acids forms **Ceramides (Cer)**, serving as the structural and metabolic hub. Cer have garnered increasing interest in their regulation of cell signaling in a myriad of biologies, including the regulation of apoptotic cell death [3, 4]. **Glycosphingolipids**, consisting of a carbohydrate moiety attached to the Cer backbone, are complex membrane components that are crucial for signal transduction, cell-cell communication, and cell differentiation [5–7]. The addition of a phosphocholine group on Cer forms **Sphingomyelins (SM)** which are among the major phospholipids found in mammalian tissues and contribute to maintaining the structure and function of the plasma membrane in cell. In much lower levels, **sphingosine-1-phosphate** (**S1P**) acts as a key signaling lipid for mammalian cells through five distinct G-Protein Coupled Receptors, promoting cell survival, cytoskeletal reorganization, and cell motility [8]. Also, in low concentrations is another signaling sphingolipid, **ceramide-1-phosphate (C1P),** which is formed by the addition of a phosphate group to the primary hydroxyl group on Cer and has been linked to pro-mitogenic and pro-inflammatory functions [9].

Understanding the intricacies of the metabolic network in biological systems necessitates the profiling and quantification of sphingolipids involved. While liquid chromatography coupled to mass spectrometry (LC–MS) is commonly employed for lipid analysis, its major limitation lies in the lack of spatial information as it measures lipids from mixtures extracted from biological samples. Positioned at the forefront of sphingolipid research [10], MALDI-MSI emerges as a novel molecular imaging technique, enabling the detection of sphingolipids directly within tissues and cell cultures and providing insight into their spatial distribution. When integrated with traditional histological techniques, such as hematoxylin and eosin (H&E) staining, MALDI-MSI discerns sphingolipid distribution across histologically distinct regions within tissues. Furthermore, recent advancements in MALDI-MSI have demonstrated enhanced spatial resolution and improved precision in single-cell imaging [11, 12], 3D imaging [13] and multimodal imaging [14] with the current emphasis often revolving around prevalent lipid species. Despite these strides in MALDI-MSI, substantial challenges persist in this technology concerning ionization, detection, and validation of lipids, especially for low abundant species.

Previous MALDI-MSI studies showed that sphingolipids can be imaged from biological samples using different combinations of MALDI parameters, including matrix selection [15–17], ion polarity mode [16, 17], mass spectrometer configuration [18, 19], as well as the laser settings. A summary of current methodologies using MALDI-MSI for imaging various sphingolipids is provided in **Table 1**. The vast majority of prior MALDI-MSI studies for sphingolipid analysis was conducted on tissue samples. In positive ion mode, 2,5-Dihydroxybenzoic acid (DHB) and α-cyano-4-hydroxycinnamic acid (CHCA) matrices have been most commonly utilized for the detection of SMs [20–26] [27–30], and ceramides [20, 21, 26, 31] [30], glycosphingolipids [32], S1P and C1P [20, 21]. A binary matrix from a mixture of DHB and CHCA has also been applied for Cer detection in positive ion mode [27]. On the other hand, 1,5-Diaminonaphthalene (DAN) matrix is frequently used in negative ion mode, with occasional studies reporting its use in positive ion mode to detect C1P [33, 34], SM [33–35], and Cers [33–36]. More recently, the 2,6-Dihydroxyacetophenone (DHA) matrix has become more common for sphingolipid imaging [37], being used in both positive and negative ion modes to image C1P[38], SMs[38–40], Cers [16, 17], and gangliosides [40, 41]. 9-Aminoacridine (9-AA) matrix has been utilized to image C1P, SM, gangliosides [42, 43], and Cers [43] in negative ion mode. Lastly, some studies have explored the use of alternative matrices, such as 3-Aminophthalhydrazide (Luminol or 3-APH), in both positive and negative ion modes to image SM, C1P, and GlcCer [44]. Considering each lipid species, SM has been most often visualized using DHB in positive ion mode, ceramide using DAN in negative ion mode, GluCer and LacCer using either DHB in positive ion mode or DAN in negative ion mode, gangliosides with DHA in negative ion mode, S1P with DHB in positive ion mode, and C1P with DAN or 9AA in negative ion mode.

**Table 1.**
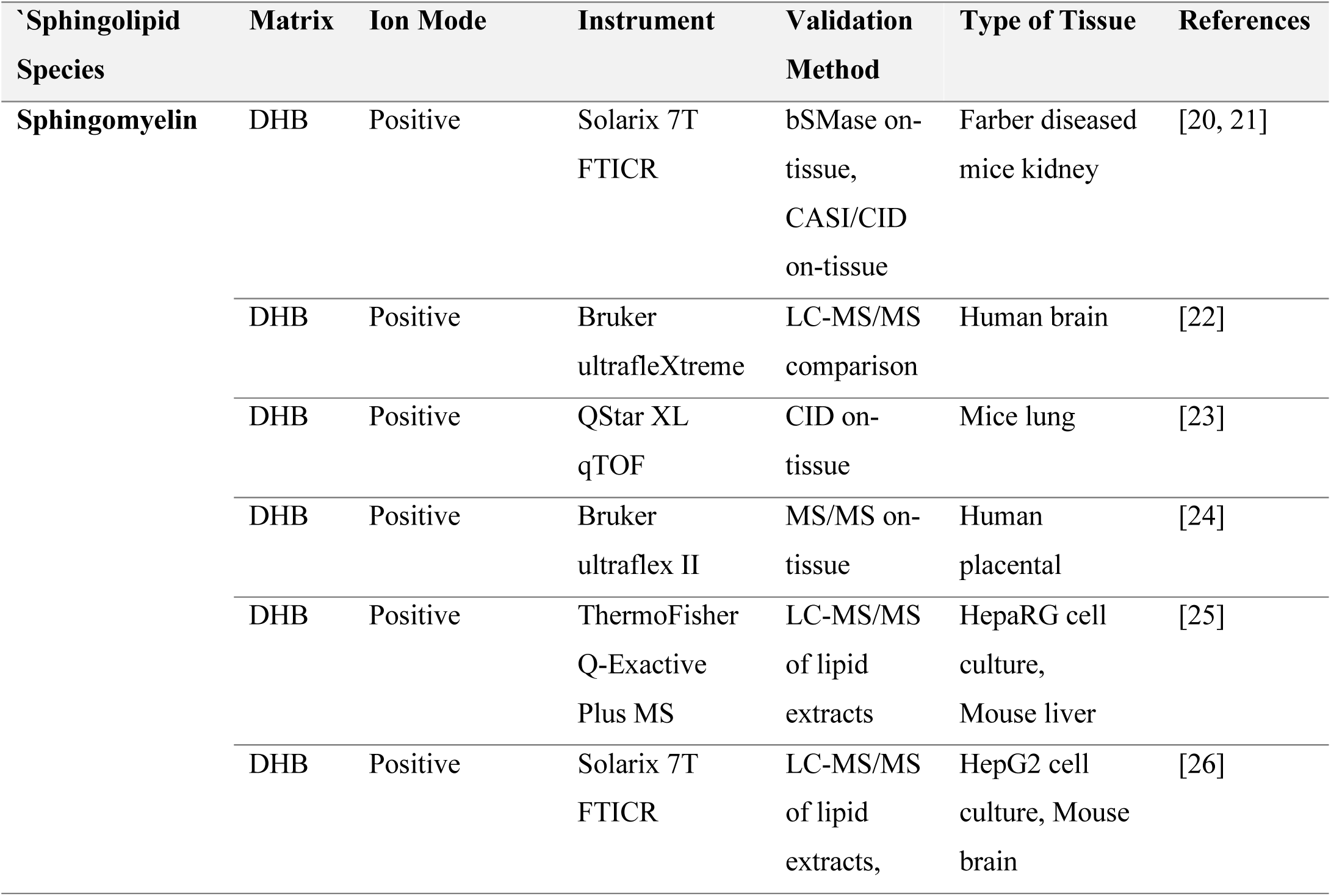
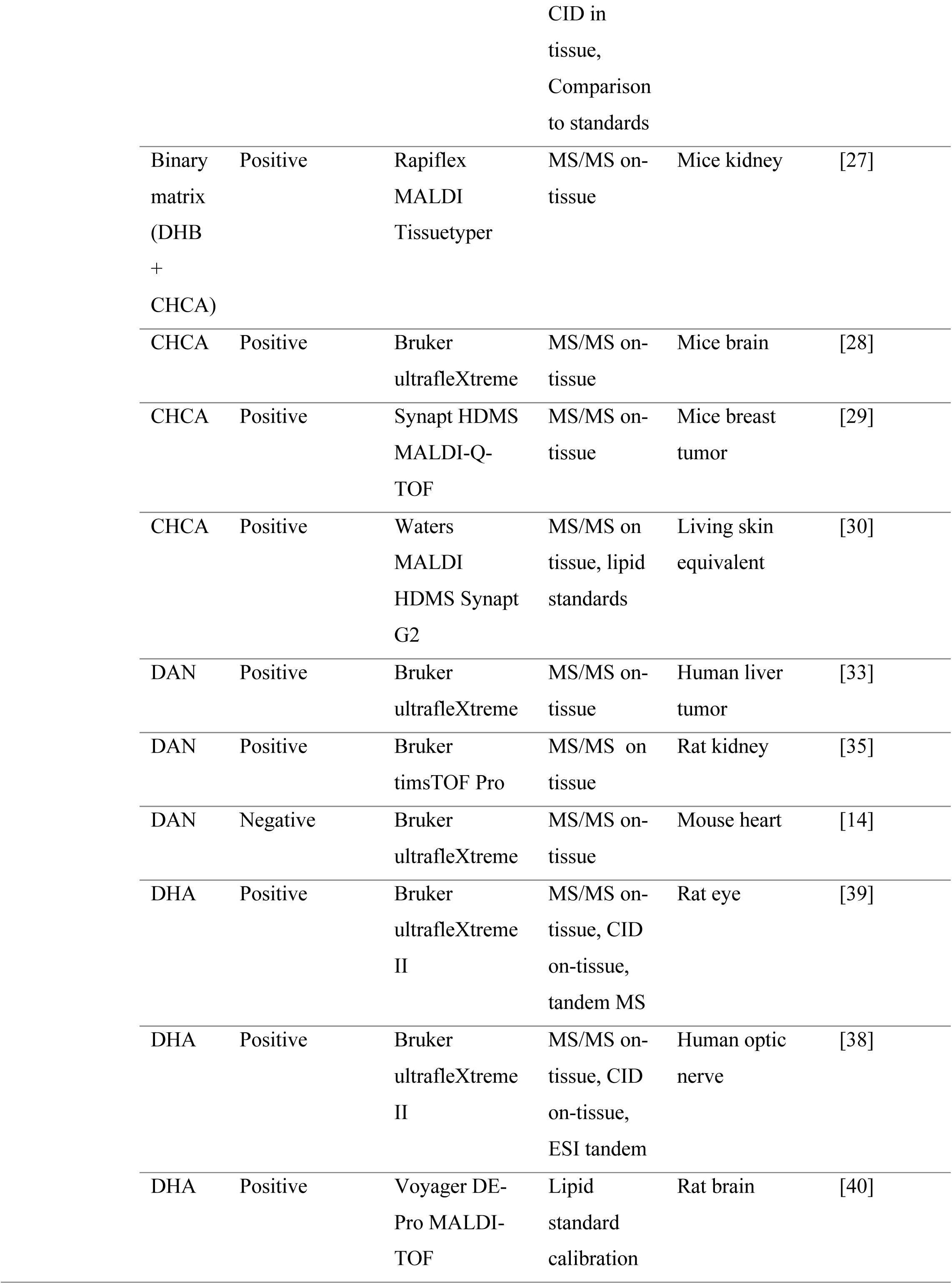
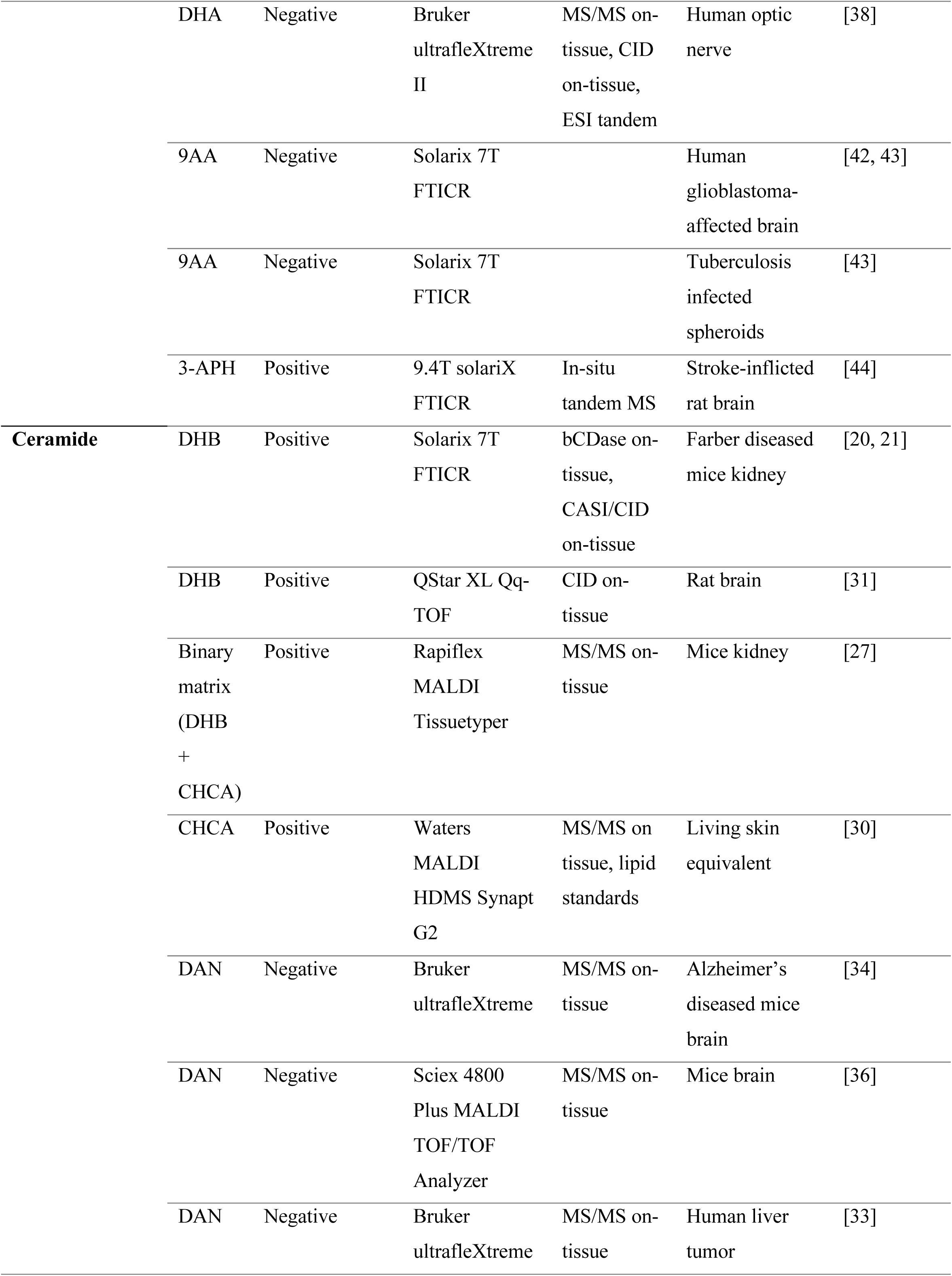
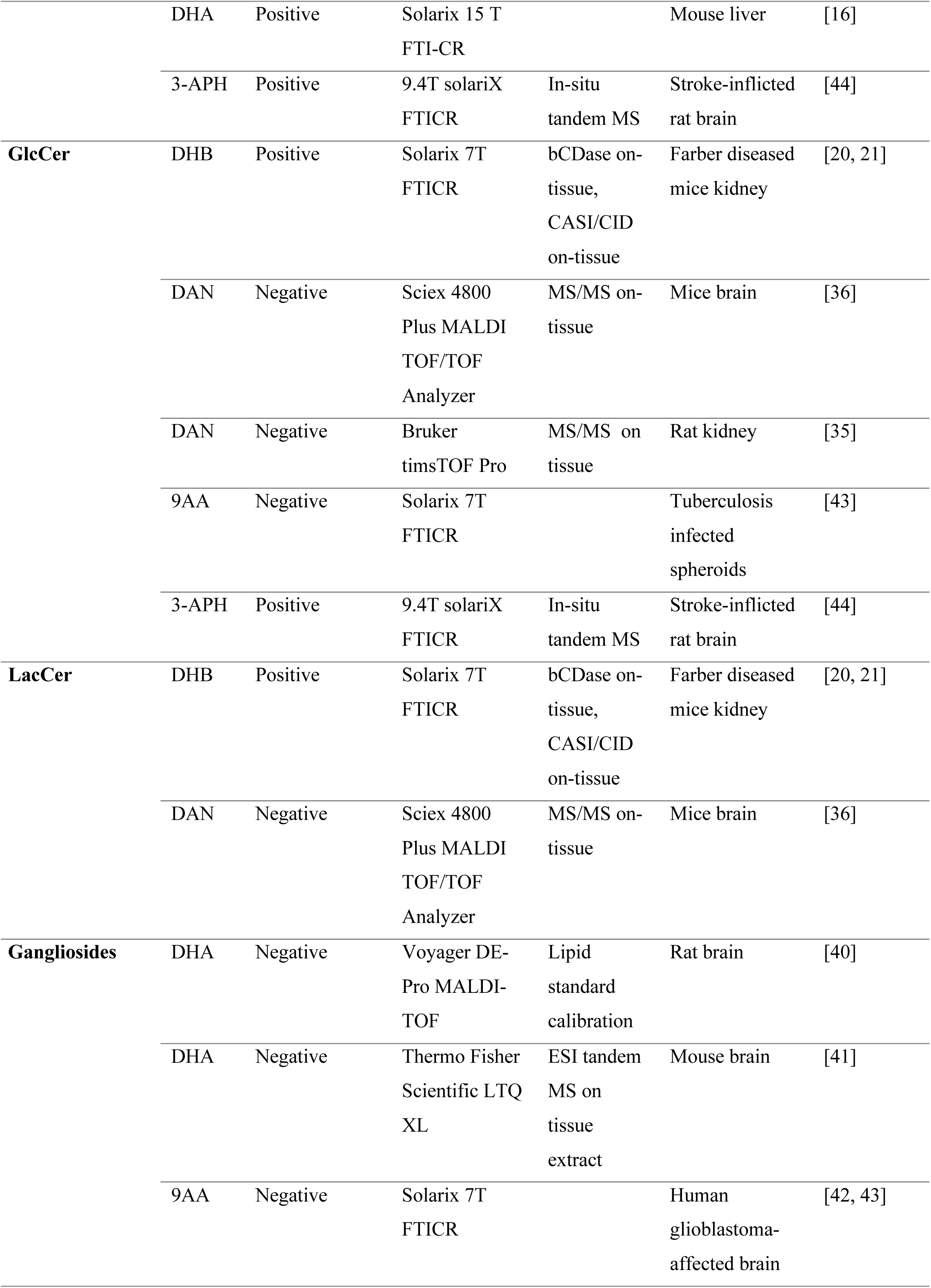
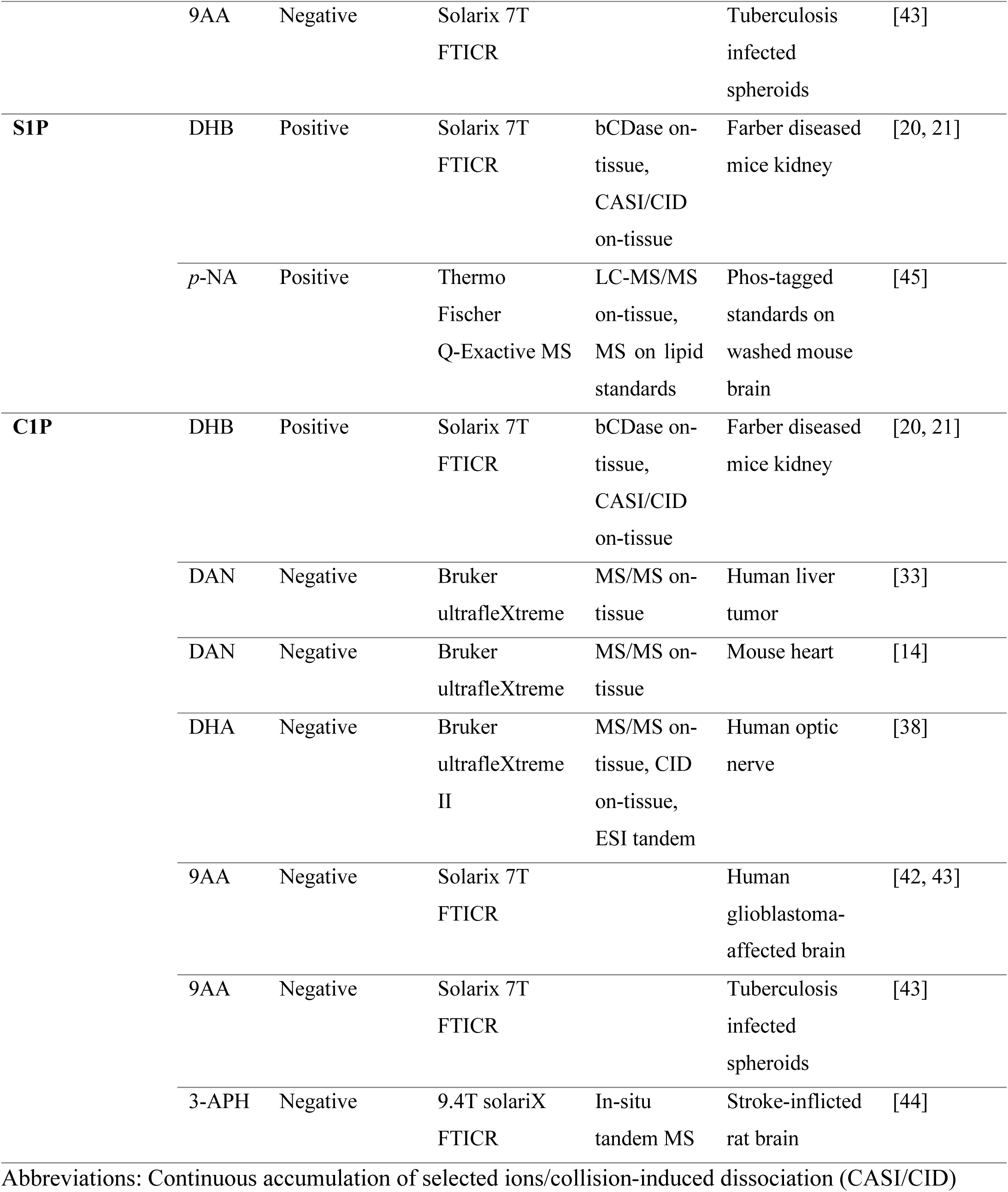
Summary of published methodologies using MALDI imaging for analysis of sphingolipid species.

Despite these recent advancements with MALDI-MSI of sphingolipid species, shortcomings/challenges still persist within the existing reported methodologies. First, using pure lipid standards, most studies do not verify the expected ionization pattern and detection under their respective experimental conditions. Second, the low level of Cers and sphingoid bases represents a challenge for their validation in biological samples by MS/MS. For example, while some publications claim the detection of S1P and C1P (very low-abundant sphingolipids, difficult to detect even by conventional LC-MS), they fail to provide convincing evidence regarding signal validation. The difficulty in obtaining strong MS-MALDI signals makes the generation of MS/MS spectra not always possible. This does not allow for the confirmation of the chemical structure of the analyte and the differentiation of it from other possible isobaric compounds. Therefore, a parallel LC-MS/MS analysis of tissue extracts is often performed to confirm the presence of certain species. To make it relevant for MALDI-MSI, local quantification of lipids can be conducted on tissue section with LC-MS/MS, provided that small tissue areas of interest are precisely collected by laser-capture microdissection [46]. However, the different ionization techniques used for electrospray ionization (ESI) and MALDI make the comparison not straightforward. A more direct approach to validate a purported lipid signal is to modulate its level by applying on the sample enzymes that can utilize the target lipid as substrate and consume it. Using this approach, SM and Cer were identified via on-tissue digestion of kidney sections with bacterial sphingomyelinase (bSMase) and ceramidase (bCDase), respectively [20, 21]. Notably, Sph, the product of Cer digestion by bCDase, was not reported in that study.

Third, the potential fragmentation of structurally complex sphingolipid species is frequently overlooked, with the danger to report artifact signals as real ones. This may constitute a complication for MALDI-MSI when it comes to detecting simpler lipid species structurally identical to the released fragments [47]. When fragments are produced by the high-abundance complex sphingolipids (glycosphingolipids and SMs), even a small percentage of fragmentation of the parent molecule (e.g. 1/5000 in the case of SM) may produce an amount of fragment that is much higher than the endogenous low-abundant species (e.g. C1P or S1P), and it can be mistaken as the signal of the natural species. Modulation of the laser intensity and choosing the proper matrix may alleviate/prevent molecule fragmentation.

Furthermore, to our knowledge, there is a lack of studies investigating Sph, an important bioactive lipid, with no published data available. Lastly, sphingolipid analysis using MALDI-MSI has been mostly conducted on tissues, with only a handful of publications utilizing cell cultures, where signals are much weaker.

These challenges underscore the need to refine MALDI-MSI methodology and delineate a more comprehensive and accurate analysis of the sphingolipid landscape. In this work, we employed commonly used matrices for the analysis of sphingolipids, namely DHA and DHB (positive ion mode) and DAN (negative ion mode). We started with analysis by MALDI-MS of reference sphingolipid standards in a ratio that approximates their relative contents found in biological samples; this allowed us to establish the extent of their ionization and to determine the impact of the generation of fragments from abundant parent molecules detected as simpler sphingolipid species, such as C1P, generally present in biological samples at a much lower concentration. Experiments in cell cultures grown on chamber slides and treated with sphingolipid enzymes and inhibitors allowed the validation of MALDI-MSI signals, differentiating them from isobaric compounds (same m/z) and excluding the artifactual effect of fragmentation of more complex species. Finally, the most appropriate experimental conditions for the various sphingolipids established upon reference standards and cell cultures were applied to murine mammary cancer tissue.

## MATERIALS AND METHODS

### Chemicals and Reagents

Methanol (LC-MS grade) (MeOH), H_2_O (LC-MS grade), and trifluoroacetic acid (TFA) (LC-MS grade) were purchased from Fisher Scientific (Waltham, MA). Acetone (liquid chromatography LiChrosolv®) and Acetonitrile (hypergrade for LC-MS LiChrosolv®) (ACN) were purchased from Sigma-Aldrich (Millipore Sigma, Burlington, MA). All lipid standards were acquired from Avanti Polar Lipids LLC (Alabaster, AL). Bacterial sphingomyelinase (bSMase) was obtained from Sigma-Aldrich (Millipore Sigma, Burlington, MA). Recombinant ceramidase from *Pseudomonas aeruginosa* (pCDase) was purified as described previously [48]. 2,5-Dihydroxybenzoic acid (DHB) matrix was from Thermo Fisher Scientific (Waltham, MA), 1,5-Diaminonaphthalene (DAN) matrix was from Millipore Sigma, Burlington, MA and 2,5-Dihydroxyacetophenon (DHA) was from Bruker Daltonics (Billerica, MA).

### Reference standards

The lipid standards were dissolved and diluted in methanol for use in experiments. The stock concentrations of the various sphingolipids reflected their approximate biological ratios found in breast cancer tissue and cells: SM:ceramide (Cer) = 100:1, Cer:glucosylceramide (GlcCer):lactosylceramide (LacCer):GM gangliosides = 1:1:1; SM:sphingosine (Sph) = 500:1, SM:S1P = 5000:1, SM:C1P = 5000:1. To obtain detectable signals from the lowest abundance species, S1P and C1P, the stock concentrations of these sphingolipids were set at 1 µM. The stock solutions of other sphingolipids followed the ratios indicated above: stock solutions for Cer, GlcCer, LacCer, GM gangliosides were at 50 µM; SM stock was at 5000 µM; and Sph stock was at 10 µM.

The samples for MALDI-MS analysis were then prepared by mixing the matrix solution with the standard solution at a 1:1 volume ratio, spotting 0.5 µL of the mixture onto a target plate. MALDI-MS spectra were collected from the replicates of reference standards on a timsTOF fleX mass spectrometer (Bruker Daltonics, Billerica, MA). Three biological replicates, with three technical replicates for each (with 10 individual spectra each), were analyzed. In some experiments, internal standards (C17-Cer and C17-Sph) were added to reference standard samples to normalize MALDI-MS signals and correct experimental variations.

### Cell culturing

HeLa (CCL-2, human adenocarcinoma epithelial cells) cells were used for cell culture experiments and were purchased from ATCC, Manassas, VA. Cells were cultured in Dulbecco’s Modified Eagle’s Medium (DMEM) (Corning Inc., NY) supplemented with D-glucose, L-glutamine, and 10% fetal bovine serum (FBS) (Gibco, Hampton, NH). Cells were maintained at 37°C and 5% CO_2_ in a humidified incubator.

For sphingolipids detection via MALDI imaging, an 8-well glass chamber Millicell EZ slide (Millipore Sigma, Burlington, MA) was used. HeLa cells were seeded at a cell density of 4 × 10^4^ cells per chamber and were grown for 24 hrs in DMEM containing 10% FBS at 37°C and 5% CO_2_. Prior to enzyme treatments, cells were serum starved for 24 hrs. Then, HeLa cells were treated with bSMase (100 mU/ml) at 37°C for 10 mins, or without bSMase (control). After bSMase treatment, cells were washed twice with PBS (Corning Inc., NY), and processed for Sph or S1P measurements. For Sph measurements by MALDI-MSI, cells were immediately fixed with 8% paraformaldehyde (PFA) (Millipore Sigma, Burlington, MA) for 5 mins at room temperature (RT) and, when appropriate (as indicated in **Figure 2**), samples were then treated with purified pCDase [48] (3.5 mg of proteins/ml) at a final concentration of 2 μl of stock enzyme solution/ml in DMEM for 1 hr at 37°C and 5% CO_2_ or without pCDase treatment (control). For S1P measurements by MALDI-MSI, the fixation step between bSMase and pCDase treatments was omitted. After the pCDase treatment, cells were processed as follows: they were washed with PBS, fixed with 8% PFA for 5 mins at RT, rinsed with PBS, washed with 150 mM ammonium formate (Millipore Sigma, Burlington, MA), and rinsed with PBS one last time. Finally, cells were dried overnight in a vacuum desiccator at RT before matrix application.

For measurements of hexosylceramides (HexCers, detected as GluCer plus GalactosylCer) and gangliosides, HeLa cells (5 × 10^5^) were seeded in 60 mm dishes in DMEM containing 10% FBS at 37°C and 5% CO_2_. After 24h, cells were washed twice with DMEM containing 1% FBS and incubated with the same medium supplemented with or without 100 nM eliglustat. Medium was changed every 48h (retaining the respective treatment conditions). After 7 days of treatment, cells were seeded in the chamber slide, grown to confluency in their respective treatment conditions (approximately for 48h), washed with PBS twice, fixed with 8% PFA for 5 minutes at RT, washed with PBS followed by 150 mM ammonium formate, rinsed with PBS one last time, and dried in a vacuum desiccator.

For sphingolipid analysis by LC-MS/MS, Hela cells (5×10^5^ cells) were plated in 60mm dishes and treated as indicated above for the MALDI glass chambers with no fixation step between bSMase and pCDase treatments. Lipid extraction and analysis was performed as previously described [49].

### Animal Model and Tissues

Breast cancers spontaneously formed in the mouse mammary tumor virus-polyoma middle tumor-antigen (MMTV-PyMT) transgenic mice were used for this study. These mammary tumors closely resemble human breast cancer tumors in terms of genomic landscape [50].

Mammary gland tumors start to develop at 12 weeks of age; at 16 weeks, mice were sacrificed, and tumors were immediately snap frozen and embedded in 5% Carboxylmethyl Cellulose (CMC) medium at −80°C. The CMC embedded freshly frozen tissues were cryosectioned at a thickness of 10 µm on a Leica CM1510S Cryostat, they were mounted on IntelliSlides (Bruker Daltonics, Billerica, MA), and kept at −80°C until analysis. Serial sections were used for MALDI-MSI and histological analysis with H&E staining.

### MALDI Mass Spectrometry Imaging

Prior to MALDI-MSI analysis, cell culture chamber slides/tissue slides were dried in a vacuum desiccator at RT for 30 mins. Slides were either sprayed with DHB or DAN matrix solution. For analysis in positive ion mode, DHB matrix solution was prepared at a concentration of 40 mg/mL in 70:30 of MeOH:water by vol and 0.1% (v/v) TFA, whereas for analysis in negative ion mode, DAN matrix solution was prepared at a concentration of 5 mg/mL in 90:10 of acetone:water by volume. Additionally, for initial optimization experiments, DHA matrix solution was also tested and prepared at a concentration of 15 mg/mL in 80:10:10 of ACN:MeOH:water by volume and 0.1% TFA (v/v) for analysis in positive ion mode. The matrix solutions were then sprayed onto each slide using an HTX M5 Sprayer (HTX Technologies, LLC, NC, USA). MALDI imaging was performed on timsTOF fleX (Bruker Daltonics, Billerica, MA) over the entire cell chamber/tissue section with 20 μm laser application. To image the same chamber/section multiple times with different methods for different lipid targets, a laser spot offset of 20 μm to a raster width of 40 μm was applied using flexImaging (Bruker Daltonics, Billerica, MA). Specific methods were used in positive or negative ion mode depending on the targeted lipid, detecting mass ranges within 200 to 2000 m/z. MSI images were generated, and the mean spectrum of MS signals was exported using SCiLS Lab software (Bruker Daltonics, Billerica, MA) for cell culture experiments or using home-made Python scripts (https://github.com/SBU-CC-MIU/MIU-MZ-to-Image) for tissue slides. Sphingolipids were annotated using MetaboScape software (Bruker Daltonics, Billerica, MA) with m/z tolerance set to 10 ppm and were cross-validated against the LIPID MAPS database.

### Validation of MALDI-MSI signals

Cell culture experiments conducted using different combinations of exogenously added bacterial sphingolipid enzymes were utilized to evaluate the reliability of the MALDI signals.

MALDI-MS/MS was performed with collision-induced dissociation (CID) to detect fragmentation patterns for SM ions. Structural assignments were made by a direct comparison between the MS/MS spectra collected from tissue and the data from reference standards.

### H&E Staining

H&E staining was performed on flash frozen tissue sections consecutive to those utilized for MALDI-MSI section.

## RESULTS

### Evaluation of DHA, DHB and DAN matrices on detection of standard sphingolipids by MALDI-MS

MALDI-MS signals corresponding to pure sphingolipid standards were evaluated as a function of matrices, selection of ion detection mode, and laser power. We applied and evaluated the most common protocols reported in the literature (DHA and DHB, and DAN matrices in positive and negative ion detection modes, respectively) on the MALDI-MS signals of reference standards (**Table 2**). Methanolic solutions of standards for Sph, SM, S1P, Cer, C1P, GlcCer, LacCer and GMs were spotted in concentrations that generally recapitulate their relative mass abundance found in tissues [51]. To allow MALDI-MS signal detection, 1μM S1P and C1P were utilized as reference concentrations for the lowest abundant species (**Table 2**). The MALDI-MS data sets were collected with the laser power set at just above the ionization threshold for each matrix (25% for DAN, 35% for DHB and 30% for DHB) to minimize fragmentation of lipid species.

**Table 2:** MALDI signal of different sphingolipids using standards mixed with DAN, DHB and DHA matrices evaluated in negative (-) or positive (+) ion detection mode. DAN signals were acquired in negative ion detection mode and 25% laser power; DHB signals were acquired in positive ion detection mode and 35% laser power; DHA signals were acquired in positive ion detection mode and 30% laser power. Highlighted in shades of green are conditions where a stronger performance is observed (darkest green for best performance). For positive ions of SM, Cer, GlcCer, and LacCer, only [M+H]^+^ (protonated) ions are included in the analysis. For positive ions of Sph, both [M+H]^+^ (protonated) and [M-H_2_O+H]^+^ (water loss) ions are considered. For negative ions of Sph, S1P, Cer, C1P, GlcCer, and LacCer, only [M-H]^-^ (deprotonated) ion is considered. Each replicate number corresponds to the average of 3 individual spots (with 8-10 point acquisition from each spot).

| Sphingolipid Standards | Concentrations tested (uM) | Replicate #1 |  |  | Replicate #2 |  |  | Replicate #3 |  |  |
| --- | --- | --- | --- | --- | --- | --- | --- | --- | --- | --- |
|  |  | DAN - | DHB + | DHA + | DAN - | DHB + | DHA + | DAN - | DHB + | DHA + |
| C16-SM | 5000 | 12294 | 11420 | 134307 | 16219 | 20124 | 144048 | 9806 | 11525 | 4746 |
| C16-Cer | 50 | 89267 | 4324 | 12477 | 23570 | 6095 | 13795 | 41002 | 1518 | 5204 |
| C16-GlcCer | 50 | 50940 | 485 | 2207 | 6954 | 669 | 4892 | 32149 | 598 | 2429 |
| C16-LacCer | 50 | 43411 | 152 | 723 | 1141 | 188 | 1153 | 54846 | 203 | 624 |
| C18-Cer | 50 | 113098 | 1043 | 9478 | 25552 | 1213 | 4770 | 49656 | 1076 | 1504 |
| C18-GlcCer | 50 | 107749 | 1636 | 1072 | 25305 | 1030 | 877 | 61763 | 643 | 1881 |
| C18-LacCer | 50 | 176979 | 142 | 860 | 9440 | 74 | 449 | 30376 | 160 | 560 |
| C18-GM1 | 50 | 64253 | 1353 | 4505 | 8466 | 274 | 729 | 5614 | 230 | 4000 |
| C18-GM3 | 50 | 53978 | 2426 | 4655 | 11653 | 317 | 5886 | 11624 | 603 | 5213 |
| Sphingosine | 10 | 809 | 22992 | 41454 | 598 | 10290 | 105257 | 417 | 9177 | 53662 |
| C16-C1P | 1 | 800 | 170 | 707 | 2362 | 1716 | 235 | 2362 | 205 | 534 |
| S1P | 1 | 730 | 841 | 771 | 153 | 756 | 915 | 151 | 387 | 397 |

A summary of the ions detected for each standard is provided in **Supplementary Table 1**. In agreement with previous publications [20–24], positive ion mode with DHB matrix produced reliable signal for SM. As indicated in **Table 2**, SM can also be detected using DAN in negative ion mode and DHA in positive ion mode.

For Cer, GlucCer, LacCer, and GMs, negative ion detection mode with DAN matrix yielded consistently higher MS signals compared to DHA or DHB in positive mode (**Table 2**), in good agreement with results from previous publications [33, 34, 36]. Ceramide imaging in positive ion mode with DHB matrix [20, 21, 31] was also previously reported. However, in our conditions, the ceramide signal acquired with DHB in positive mode produced a qualitatively weaker signal than the DAN matrix in negative mode (**Table 2**).

Among the less abundant species, Sph presented a stronger signal with DHA and DHB matrices in positive ion mode compared to than with DAN (**Table 2**). In DHB, the water loss peak [M-H_2_O+H]^+^ at

### 282.279 m/z, typically used to identify Sph in LC-MS, was dominant compared to the protonated peak [M+H]^+^ at 300.290 m/z

While still low, the S1P signal was detectable with all three methods, with a slightly better performance observed using DHB and DHA in positive ion mode (**Table 2**).

As shown in **Table 2**, C1P was ionizable by DAN matrix, and it was best detected in negative ion mode. One recurring challenge with these experiments was the variability of the signals among biological replicates and, for each biological replicate, within its technical replicates. We sought to establish whether including normalization with an internal standard (IS), distinct from the authentic standards, but close in structure (added in the mixture with the matrix), would ameliorate this variability. C17-Cer (an unnatural ceramide with a C17 sphingoid backbone) was utilized as IS in DAN negative ion mode, given the good performance observed for Cer authentic standards with this method (**Table 2**). C17-Sph was utilized in DHB positive ion mode for the same reason. While still somewhat variable, results using the signal of the IS to normalize the signal intensity of each analyte show a tighter profile among the three replicates (**Supplementary Table 2**). Differently from the results in **Table 2**, here signals relative to the same authentic standard cannot be compared between DAN and DHB since two different ISs were utilized.

### Evaluation of MALDI-induced fragmentation of complex sphingolipids

After establishing the protocols that performed best for each of these sphingolipids, we investigated the potential fragmentation of the more complex sphingolipid species, a process that, in a biological sample, could interfere with the detection of simpler species, generally present at much lower amounts. **Table 3** represents the fragmentation degree of SM, Cer, GlcCer, LacCer, and GM gangliosides species tested with either DAN, DHA, or DHB matrices. The ratio of MS signals between fragment species (S1P, Cer, C1P, GlcCer, and LacCer) (**Table 3**) and their parent species reported in **Table 2** were used to quantify the degree of fragmentation. Pure standards carrying C16:0 fatty acid were used to test fragmentation of SM, Cer, GlcCer, and LacCer, while C18:0 species were used to test fragmentation of glycosphingolipids (GlcCer, LacCer, and GM) as pure C16:0-GM standards were not commercially available.

**Table 3:**
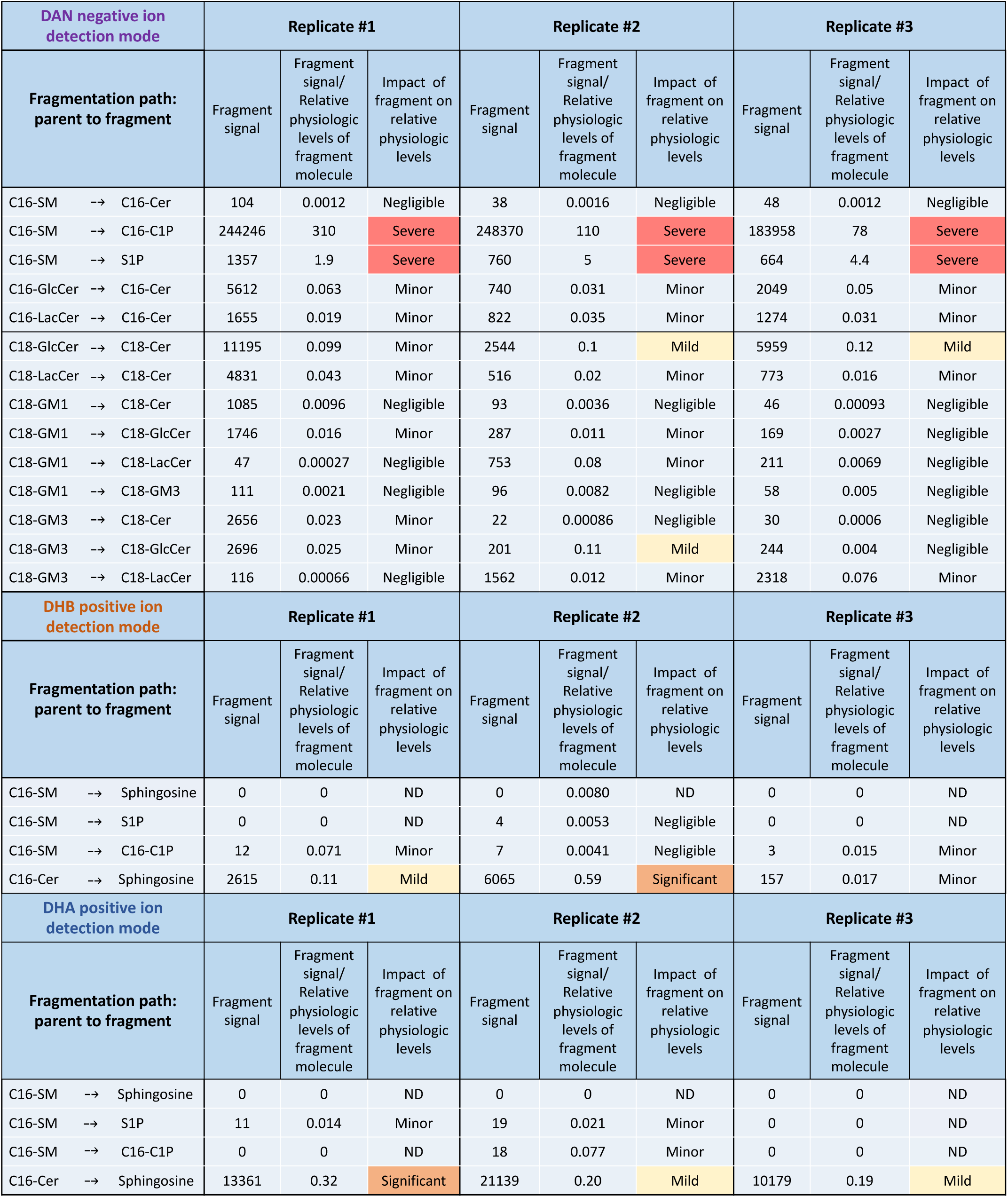
Assessing fragmentation of authentic standards and the formation of fragments that could ultimately interfere with MALDI measurements of low abundance sphingolipids. Ratios between the signal intensity of fragments produced from complex sphingolipids and the relative abundance of the same low abundance molecules in biological samples were derived using values from the fragment signal column reported in this Table and signal values of authentic standards in Table 2. Table 2 and this table report data collected in the same experiments and therefore comparable. DAN signals were acquired in negative ion detection mode and 25% laser power; DHB signals were acquired in positive ion detection mode and 35% laser power; DHA signals were acquired in positive ion detection mode and 30% laser power. Color scale red to yellow indicates high to low fragment production (ratio <0.1 indicates that production of fragment has a Minor impact; ratio 0.1-0.2 indicates that production of fragment has a Mild impact; ratio >0.2 and <0.6 indicates that production of fragment has a Significant impact; ratio >0.6 indicates that production of fragment has a Severe impact). For very low abundance signals, background matrix signal is subtracted. For positive ions of SM, Cer, GlcCer, and LacCer, only the [M+H]^+^ protonated ion is included in the analysis. For positive ions of Sph, both protonated ion [M+H]^+^ and water loss ion [M-H2O+H]^+^ are considered. For negative ions of Sph, S1P, Cer, C1P, GlcCer, and LacCer, only the deprotonated ion [M-H]^-^ is considered.

Cer can be theoretically formed by fragmentation of SM, GlcCer, LacCer, GM1, and GM3. Since Cer is best measured with DAN in negative ion mode (**Table 2**), the potential fragmentation of complex sphingolipids into Cer in this condition is most relevant. As shown in **Table 3**, no significant breakdown of complex sphingolipids into Cer was observed in all cases except for a mild/minor fragmentation of GlucCer. Also, no consistent fragmentation of GM1 and GM3 to produce GlcCer and LacCer was observed with DAN in negative ion mode (**Table 3**), which was the better performing method for measuring GlcCer and LacCer (**Table 2**). On the other hand, mild to severe fragmentation of LacCer-to-Cer and GlcCer-to-Cer and of complex glycosphingolipids into LacCer and GlcCer was observed with the DHB and DHA matrices in positive mode (**Supplementary Table 3)**. Altogether, these results suggest that, in DAN in negative ion mode, fragmentation would not substantially contribute to the MS signals for Cers, GluCer, and LacCer detected.

As shown in **Table 2**, Sph was best detected in DHA and DHB matrix and positive ion mode; under these conditions, the formation of a Sph fragment from C16-SM is negligible while fragmentation of C16-Cer can produce Sph fragments in amounts that may have a mild to significant impact (**Table 3**).

As shown in **Table 2**, S1P was best detected using DHB or DHA in positive ion mode, and in these conditions, no significant fragmentation of SM-to-S1P was observed (**Table 3**). On the other hand, severe fragmentation of C16-SM caused the formation of S1P when tested with DAN in negative ion mode, indicating that detection of S1P should be exclusively carried out in DHB or DHA in positive mode.

C1P was best detected with DAN in negative ion mode (**Table 2**); however, massive SM-to-C1P fragmentation was observed in this condition, in agreement with a previous study [47]. Considering the much larger amount of SM compared to C1P in biological samples (thousands of fold), the amount of C1P fragments formed from SM using DAN in negative mode (**Table 3**) may mask any authentic C1P signal. Results in **Table 2** suggest that a modest C1P signal may be retrieved using DHA and/or DHB in positive ion mode.

In summary, the most abundant sphingolipids can be ionized and detected effectively with a combination of DHA/DHB positive ion mode or DAN negative ion mode. For the less abundant S1P and C1P, results using DAN matrix in negative mode raise a serious warning regarding the influence of SM fragmentation on their detection. This artifact could be circumvented by using DHB (or DHA) in positive mode, although, in the case of C1P, at a cost for signal. A summary of the fragmentation patterns is provided in **Figure 1**.

**Figure 1.**
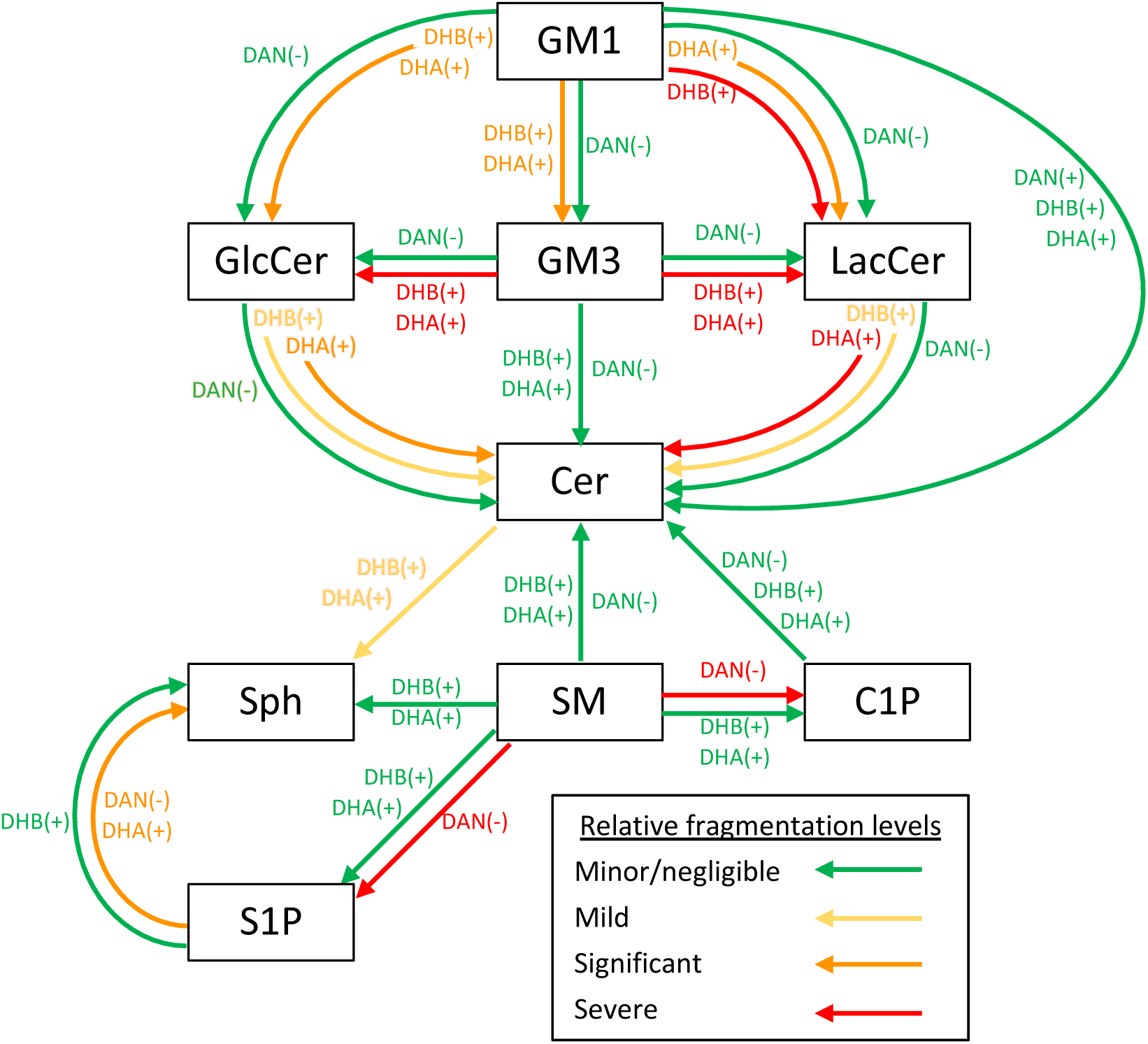
Schematic illustrating fragmentation paths of the various sphingolipid standards using DAN in negative mode versus DHB and DHA in positive mode. Colored arrows represent the relative fragmentation levels of sphingolipid standards in the indicated matrix. Assignments of relative fragmentation were made according to fragmentation values in Table 3 and Supplementary Table 3.

**Figure 2.**
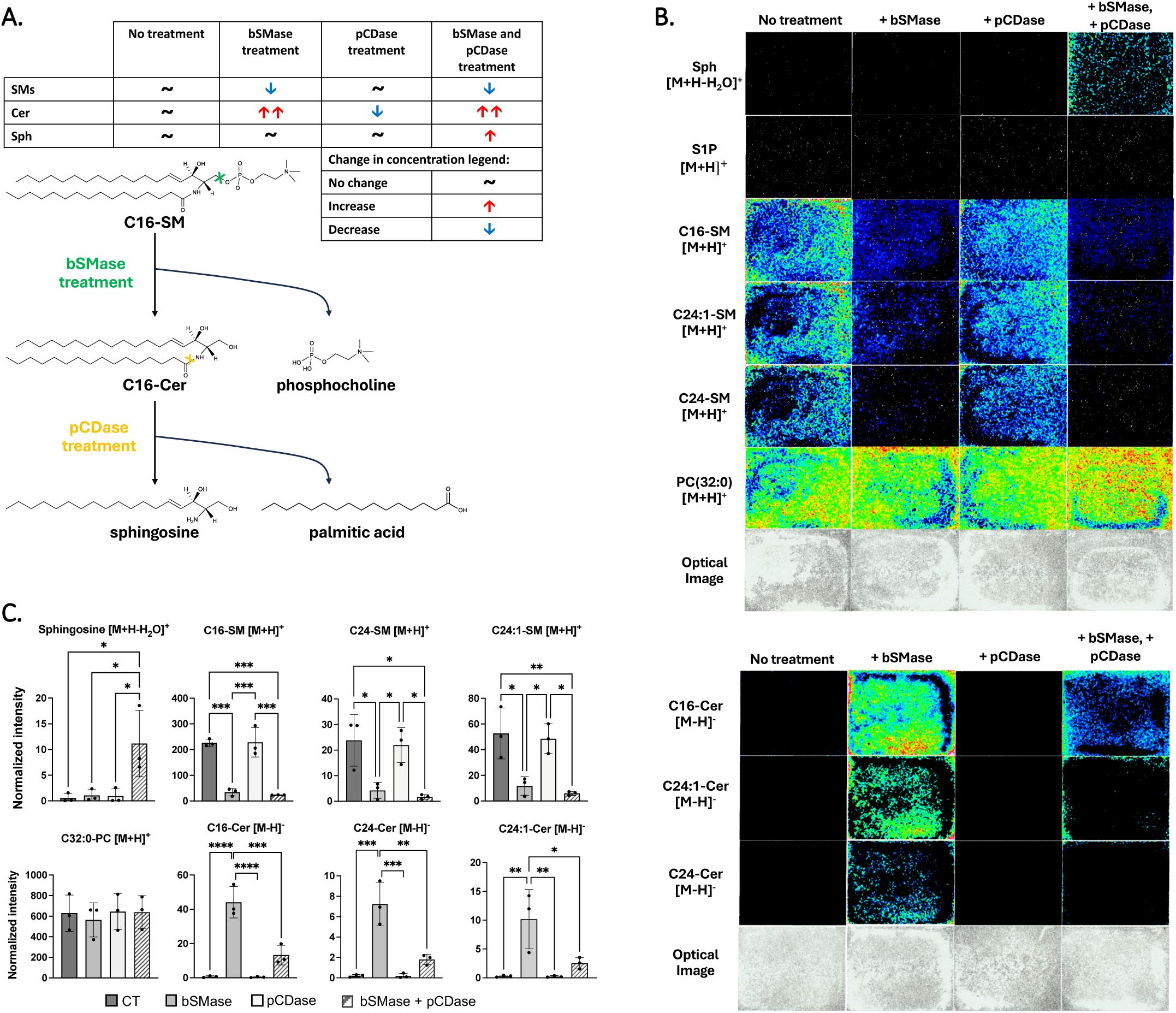
Validation of MALDI-MSI signals for SM, Cer, and Sph in cell culture. (**A**) Schematics of the reactions catalyzed by bSMase and pCDase. Treatment with bSMase (100uM/ml) cleaves SM to produce Cer and phosphocholine. pCDase cleaves Cer into Sph and fatty acid. Point of cleavage is denoted with “X”. Incubation with pCDase: 2 μl of stock solution/ml of DMEM for 1 hr; pCDase stock solution: 3.5 mg of proteins/ml. (**B**) MALDI-MSI images from HeLa cells grown and treated in chamber slides. Top panels show signals for Sph as [M-H_2_O+H]^+^ and S1P, C16-SM, C24:1-SM, C24-SM and PC(32:0) as [M+H]^+^ collected in DHB positive ion mode. Bottom panels show signals for C16-Cer, C24:1-Cer, and C24-Cer as [M-H]^-^ collected in DAN negative ion mode. Optical images are provided alongside to show cell density. (**C**) Signal intensities of sphingolipid species detected by MALDI-MSI were normalized using the optical density of the chambers as a proxy for cell number. Values represent standard deviation from the mean intensity of 3 independent biological replicates. Statistics: One-way ANOVA with post hoc Tukey’s multiple comparisons test ∗P<0.05; ∗∗P<0.01; ∗∗∗P<0.001; ∗∗∗∗P<0.0001.

### Validation of MALDI-MSI signals for SM, Cer, and Sph in cell culture

Following the MALDI-MS study on reference standards, we conducted MALDI-MSI analysis on cell cultures to validate the putative observed signals. As biological samples are a complex mix of a myriad of molecules, in addition to fragmentation, signals could also be due to the presence of isobaric compounds, and further validation should be sought. However, cell cultures are amenable to modulation of sphingolipid levels, and therefore, they can easily be used to confirm the identity of a detected signal. Furthermore, studies in cell culture would reveal the limit of MALDI-MSI detection.

To this aim, we employed treatment with exogenously applied sphingolipid modulating enzymes, bSMase, and pCDase [20, 21] and eliglustat, a highly specific and potent inhibitor of GlcCer synthase [52]. Specifically, bSMase treatment cleaves SM and produces Cer and phosphocholine, while pCDase treatment hydrolyses Cer, producing Sph and fatty acid (**Figure 2A**).

MALDI-MSI signals are shown in **Figure 2B**, and quantitation of the MALDI-MSI signal over cell density from the optical images is shown in panel **2C**. Signals for 16:0-, 24:0-, 24:1-SMs in DHB positive ion mode were readily detected in control cells, while they significantly diminished after bSMase treatment (**Figure 2B-C**). A profound increase of 16:0-, 24:0, 24:1-Cers was also observed after bSMase treatment in DAN negative ion mode, as expected (**Figure 2B**). These results were in line with substantial SM hydrolysis and Cer formation following bSMase treatment as measured by LC-MS/MS (**Supplementary** Figure 1). Cells treated with both bSMase and pCDase showed decreased Cer MALDI-MSI signal and increased Sph (DHB positive mode) (**Figure 2B-C**). Notably, the fragmentation of C16-Cer to Sph observed using pure standards (**Table 3**) was minimal, if at all present, in cells. In fact, Sph signal was scarcely detectable under bSMase treatment (**Figure 2B-C**), while Cers greatly accumulated due to the hydrolysis of SMs (**Figure 2B-C** and **Supplementary** Figure 1), and Sph was only observable in case of bSMase and pCDase (**Figure 2B-C**), where Cers levels were considerably lower and Sph level accumulated (**Supplementary** Figure 1). Of note, both Cers and Sph signals were not detectable in untreated cells.

Importantly, the enzymatic treatments did not affect the MALDI-MSI signal for phosphatidylcholine (PC), a typical isobaric compound for SM, further strengthening the overall specificity of the results. While pure standard S1P could be detected by MALDI-MS in DHB positive ion mode (**Table 2**), no S1P signal in all treatment conditions could be picked up by MALDI-MSI (**Figure 2B)**; this indicates that in cells, even in the presence of large S1P accumulation, as in case of double bSMase/pCDase treatments (**Supplementary** Figure 1), under controlled experimental conditions, cellular levels of S1P are below the limit of detection for MALDI-MSI.

Results using standards reported in **Table 3** indicated extensive fragmentation of C16-SM to form C16-C1P in DAN negative ion mode (**Figure 3A**). These results were confirmed when analyzing cells after bSMase treatment (**Figure 3B**). In fact, the deprotonated C16-C1P ion signal detected by MALDI-MSI was dependent on the presence of protonated C16-SM ions, being clearly visible in control cells and disappearing upon SM hydrolysis (bSMase treatment) (**Figure 3C**).

**Figure 3.**
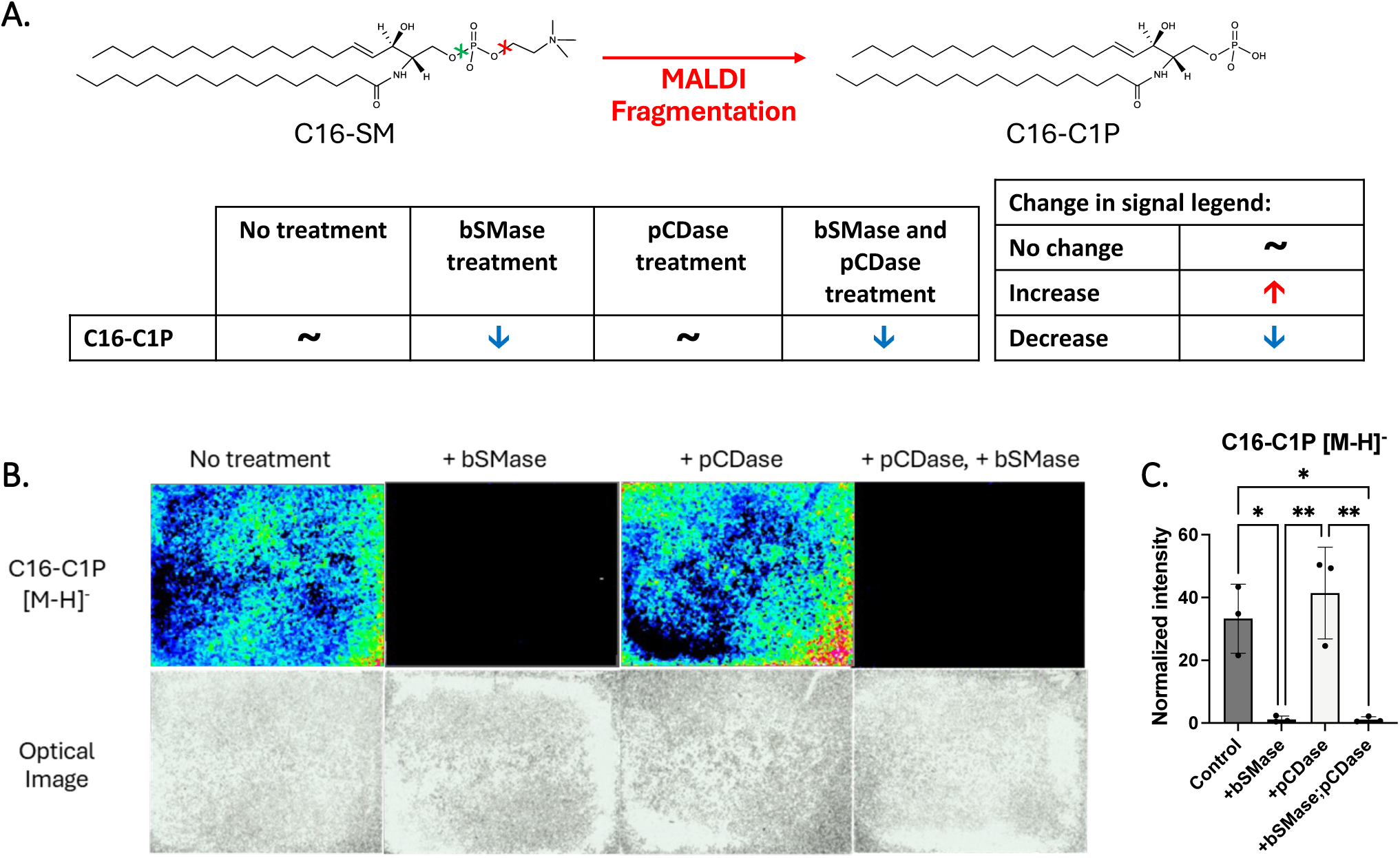
MALDI-MSI signals for C1P in cell culture using DAN negative ion mode. (**A**) Potential C16-SM fragmentation and formation of C16-C1P caused by MALDI-MSI using DAN matrix in negative ion mode. (**B**) MALDI-MSI images from HeLa cells grown and treated in chamber slides, as indicated. Panels show signals for C16-C1P [M-H]^-^ collected in DAN negative ion mode and following treatments with bSMase and/or pCDase. Optical images are provided alongside to show cell density. (**C**) Signal intensities of C16-C1P [M-H]^-^ detected by MALDI-MSI in cell culture triplicates from the various treatments were normalized using the optical density of the chambers as a proxy for cell number. Values represent standard deviation from the mean intensity of 3 independent biological replicates. Statistics: One-way ANOVA with post hoc Tukey’s multiple comparisons test ∗P<0.02; ∗∗P<0.003.

### Validation of MALDI-MSI signals for glycosphingolipids in cell culture

To assess the specificity and sensitivity of MALDI-MSI signals associated with glycosphingolipids in cells, HeLa cells were grown for a week with or without eliglustat (100nM) and then plated into chamber slides for MALDI-MSI analysis (**Figure 4A**). Eliglustat is a specific inhibitor of GlcCer synthase, therefore it blocks the main metabolic branch for the formation of glycosphingolipids (**Figure 4A**). MALDI-MSI images of Gb3, GM3, GM2, and GM1 are shown in **Figure 4B**.

**Figure 4.**
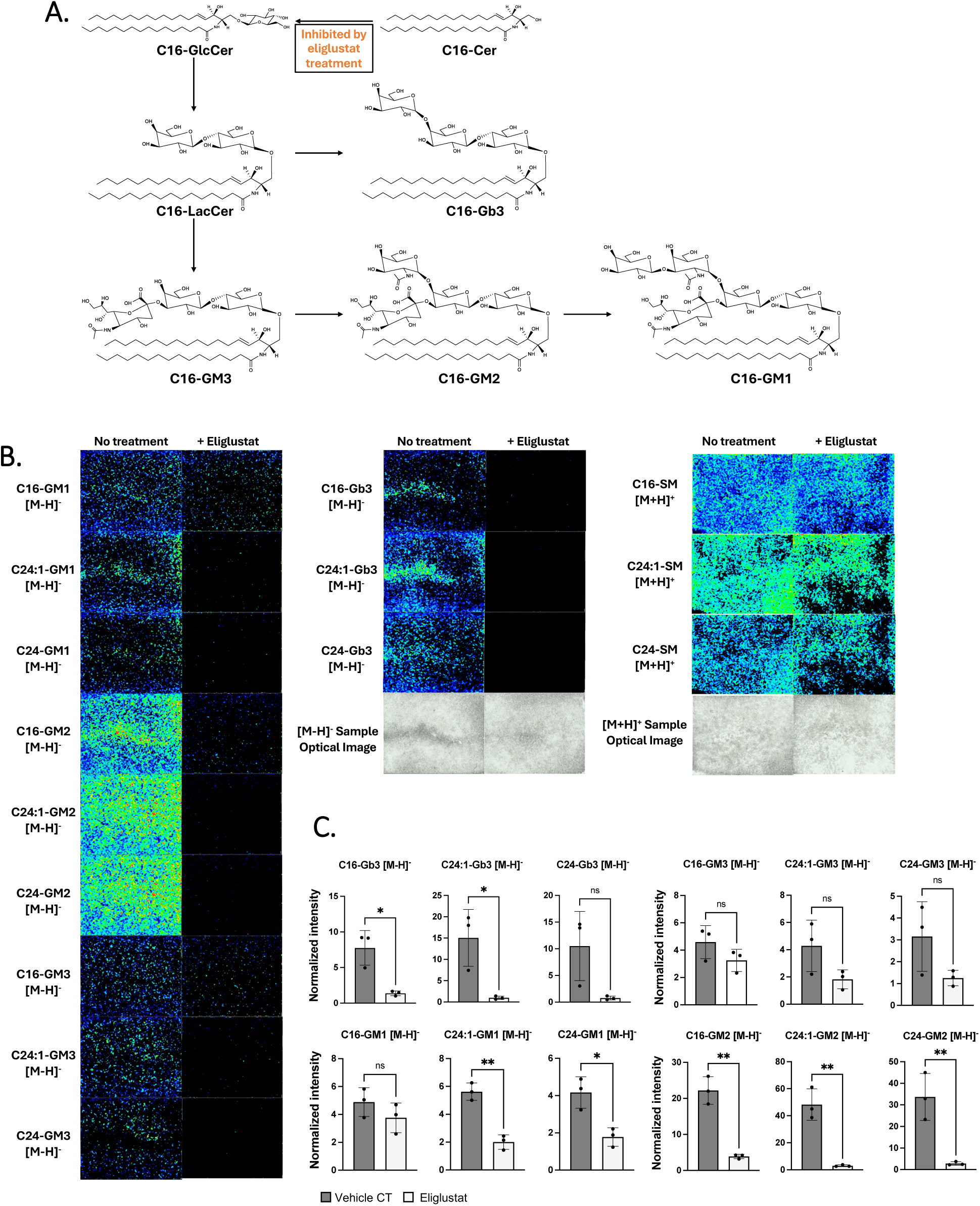
Validation of MALDI-MSI signals for glycosphingolipids in cell culture. (**A**) Eliglustat inhibits GlcCer Synthase, the enzyme that forms GlcCer from Cer. GlcCer is then used as substrate for synthesis of LacCer, a precursor to the formation of Gb3 and GMs. (**B**) MALDI-MSI images from HeLa cells grown and treated in chamber slides, as indicated. Panels show signals for [M-H]^-^ of the various glycosphingolipids collected in DAN negative ion mode after treatments with or without eliglustat. Optical images are provided alongside to show cell density. (**C**) MALDI-MSI signal intensities in cell culture triplicates from the various treatments were normalized using the optical density of the chambers as a proxy for cell number. Values represent the standard deviation from the mean intensity of 3 independent biological replicates. Statistics: Unpaired two-tailed T-test ∗P<0.03; ∗∗P<0.01.

Treatment of HeLa cells with eliglustat caused a marked reduction in various molecular species of Gb3, GM3, GM2, and GM1 MALDI-MSI signals, compared to untreated cells (**Figure 4B**). This is in line with eliglustat inhibiting GlcCer synthase, the enzyme regulating the initial step of biosynthesis for these complex glycosphingolipids. Quantitation of the signal intensities of the various glycosphingolipids detected by the MALDI-MSI and normalized by cell density (**Figure 4C**) confirmed the reduction of Gb3 and GM species.

### MALDI-MSI analysis of tissue sections using optimal experimental parameters set by MALDI-MS of standards and validation in cells

MALDI-MSI was performed on tissues, implementing the optimized parameters set using reference standards and cell cultures (**Figure 5**). Serial tissue sections from PyMT mammary tumors, flash frozen and embedded in CMC, were coated with either DHB matrix for detection of SM and Sph or DAN matrix for detection of Cer and glycosphingolipids.

**Figure 5.**
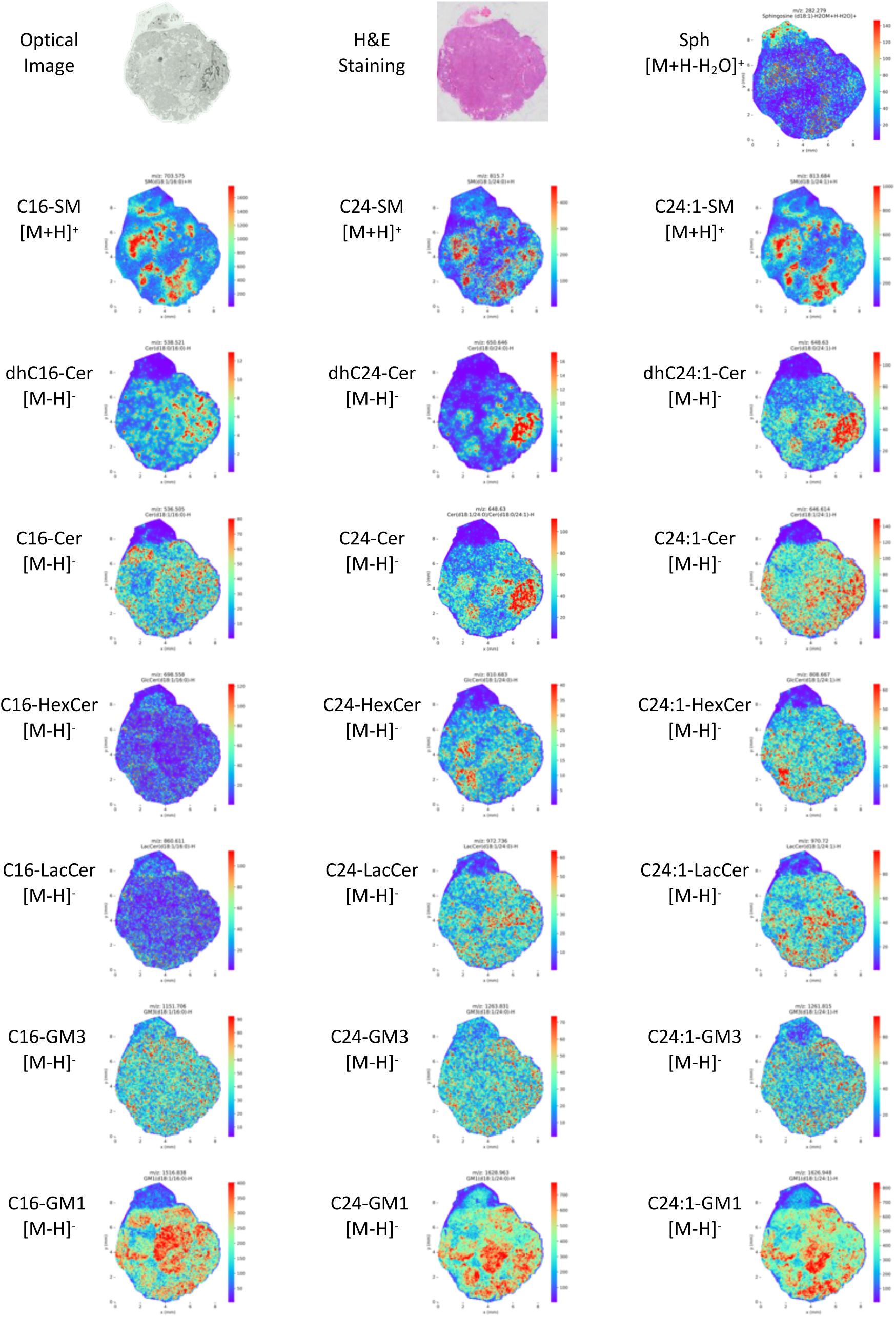
MALDI-MSI signals of major SMs, Cers, dihydroCers, Sph, hexosylCers, LacCers, and gangliosides in PyMT mammary tumor. Serial sections were fixed and analyzed for MALDI-MSI or H&E staining. MALDI-MSI sections were sprayed with DHB or DAN and signals captured in positive or negative ion detection mode, respectively. MALDI-MSI images of Sph and SMs (upper rows) were collected with DHB in positive ion mode. MALDI-MSI images of Cers, HexCers, LacCers, dihydroCers, and gangliosides were taken with DAN in negative ion mode from a consecutive section.

Figure 5 shows MALDI-MSI images of major SMs, Cers, HexosylCers (HexCers; detected as GlcCer plus GalactosylCer), LacCers, dihydroCers, gangliosides, and Sph detected from serial sections of PyMT mammary tumor. Selected ions are the same as those indicated in **Supplementary Table 1** for reference standards.

For SM, common adduct ions, [M+H]^+^, [M+Na]^+^, and [M+K]^+^ were also detected and are provided in **Supplementary** Figure 2. The SM species detected in MALDI-MSI were validated directly in tissue by MALDI-TOF MS/MS and compared to reference standards as shown in **Supplementary** Figure 3 (on-tissue fragmentation of [SM(d18:1/16:0)+H]^+^ via MALDI-MS/MS compared to the MS/MS fragmentation of the C16-SM). The fragment peak at 184.073 m/z is a characteristic ion due to the loss of the phosphocholine head group, and the peak at 264.267 m/z corresponds to [M-2H_2_O+H]^+^ ion for the sphingosine backbone.

Based on the analysis of the Sph (d18:1) reference standard by MALDI-MS, the water loss peak [M-H_2_O+H]^+^ ion at 282.279 m/z is reported from the same tissue section analyzed for SM (DHB positive mode). Interestingly, the distribution of Sph partially correlates with SM in the tissue (Figure 5).

MALDI-MSI images of Cers, dihydroCers, HexCer and LacCers were obtained in negative ion mode with DAN matrix. HexCer signal is the sum of both GlcCer and GalactosylCer ions, as these two isomers cannot be distinguished by MALDI-MSI based on m/z identification alone. However, in most tissues, the levels of GlcCer are far more abundant than those of GalCer. Of note, Cer profiles are distinct from those of SMs, and in some cases, they seem mutually exclusive. Of note, Cer distribution is distinct from Sph’s suggesting that, similarly to cells, also in the PyMT, Cer fragmentation into Sph is not prevalent. Noticeable were the distinct distribution patterns among Cer species and glycosphingolipid species.

No signal was detected for S1P in DHB positive ion mode (data not shown), while a weak signal was detected in DAN negative mode (**Supplementary** Figure 4). In DAN negative mode, SM reference standards showed severe fragmentation into S1P (**Table 3**), while in cells, potential SM-to-S1P fragments were not detected. In the tissue, Pearson correlations analysis of sphingolipid species shows a poor correlation between S1P putative signal and SM (**Supplementary** Figure 5, S1P in red), suggesting that SM fragmentation did not contribute significantly to S1P m/z signal and that the putative S1P signal may represent an unidentified isobaric compound present in the section.

While no signal was detected for C1P in DHB positive ion mode (data not shown), a strong C1P signal was detected in DAN negative mode (**Supplementary** Figure 4). In this condition, massive fragmentation of SM into C1P was observed with authentic standards (**Table 2**) and in cells (Figure 3); in line with this, a correlation can be observed between the distribution of C1P and SM signals in the PyMT tissue in DAN (-) (**Supplementary** Figure 4). These results suggest that, in tissues, C1P signal in DAN negative mode may result from SM fragmentation.

## DISCUSSION

This work contributes three important deliverables for the study of sphingolipids by MALDI-MSI: 1. A logical, rigorous, and comprehensive blueprint for validated analysis of major sphingolipids; 2. A method to detect Sph, a major bioactive sphingolipid for which, thus far, a MALDI-MSI protocol has not been reported in the literature; and 3. The conclusion is that the C1P and S1P MALDI-MSI signals detected in cells and tissues with DAN negative mode are not authentic. The C1P signal in cells and PyMT tissue is the result of SM fragmentation, and the S1P signal in PyMT likely comes from the detection of an isobaric molecule.

In our study, optimal MALDI-MS parameters for sphingolipids were initially determined through the analysis of reference standards, supporting the use of DHB in positive ion mode for measurements of SM, Sph, C1P, and S1P, and DAN in negative ion mode for Cers and glycosphingolipids. Subsequently, these parameters were validated in MALDI-MSI using cell cultures. Here, SM and complex glycosphingolipids were readily detectable in basal conditions, while Cer and Sph only upon induction. S1P was not detectable, even upon great induction, and C1P was not detectable in DHB, while a false positive signal was present in DAN resulting from SM fragmentation. Finally, using the most appropriate parameters, MALDI-MSI was conducted on tissue sections (**Table 4**).

**Table 4.**
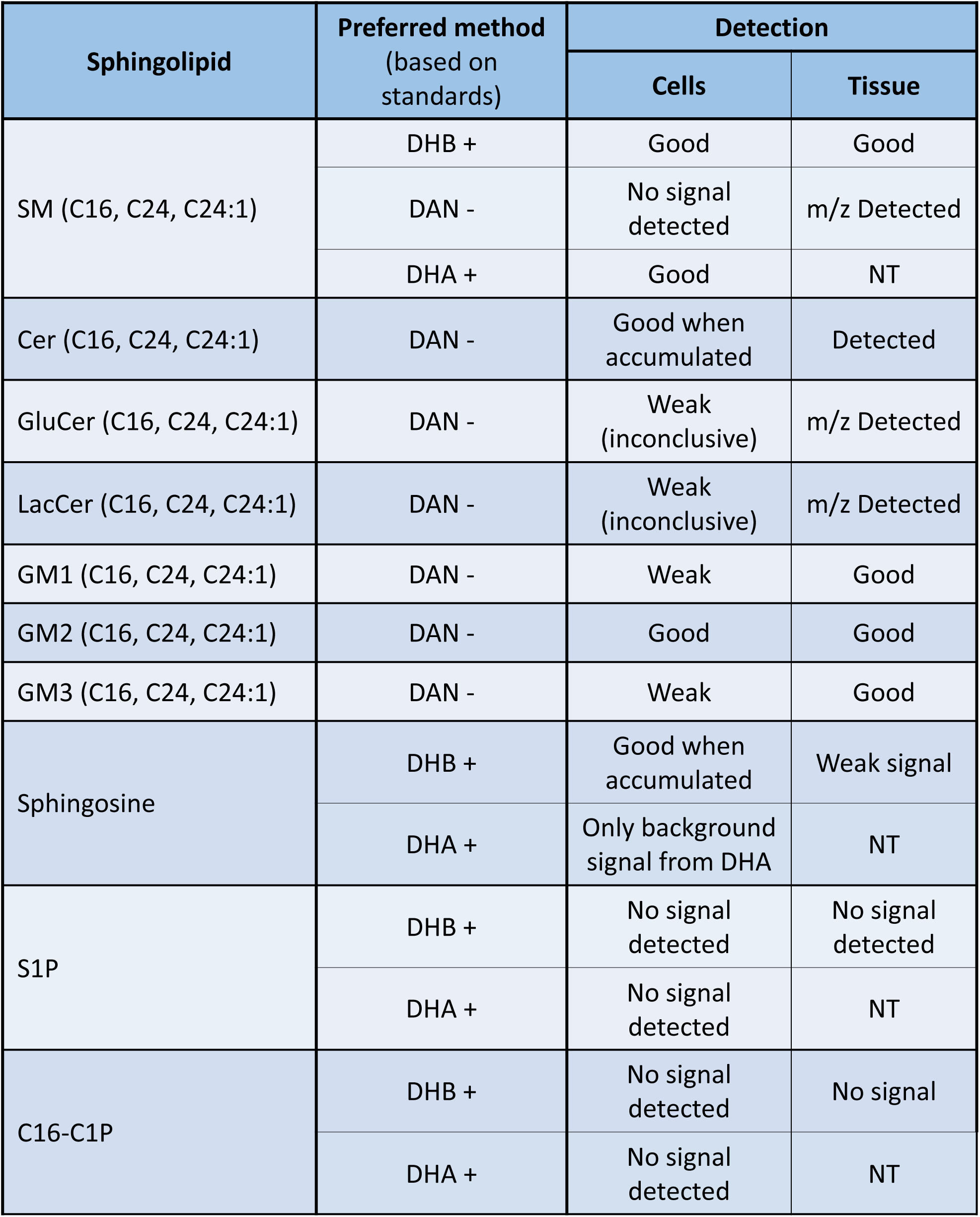
Summary of tested/validated protocols in cells and tissue. MALDI results using the better performing protocols for each sphingolipid as evaluated using authentic standards (Figure 1) are summarized in their performance in cell culture and PyMT breast cancer tissues. NT: not tested. Negative sign (–) represents negative ion mode; the positive sign (+) represents positive ion mode.

In the era of metabolomics and (pre)clinical advances, MALDI-MSI is a powerful analytical tool; however, for meaningful results, it requires rigorous validation. In this study, we have adopted different strategies to improve the validation of MALDI-MSI signals and confirm measurements in biological samples. The main advancements over the currently reported protocols are summarized in Figure 6.

1. *We have advanced the use of authentic standards by testing the addition of an IS for more consistent quantitation of the response and by studying their behavior at concentrations that reflect their relative ratios in biological samples.* Reference standards were used: i) to check the ionizability and detectability of lipid species in MALDI-MS [26, 40]; ii) for validating sphingolipid species through the comparison of their fragmentation patterns with those acquired from tissue samples in MS/MS [20, 30]; and iii) for calibration and quantitation in MALDI-MSI studies of tissue samples.[53]. However, there are technical challenges when utilizing reference standards in MALDI-MS [54]. When mixed with matrices in solutions, standards present uneven spotting, resulting in rough spot surfaces, nonuniform spot density and variable signal between repeated spots. We found that the inclusion of an appropriate IS together with the authentic standards helps to normalize this variability (**Supplementary Table 2**).

**Figure 6.**
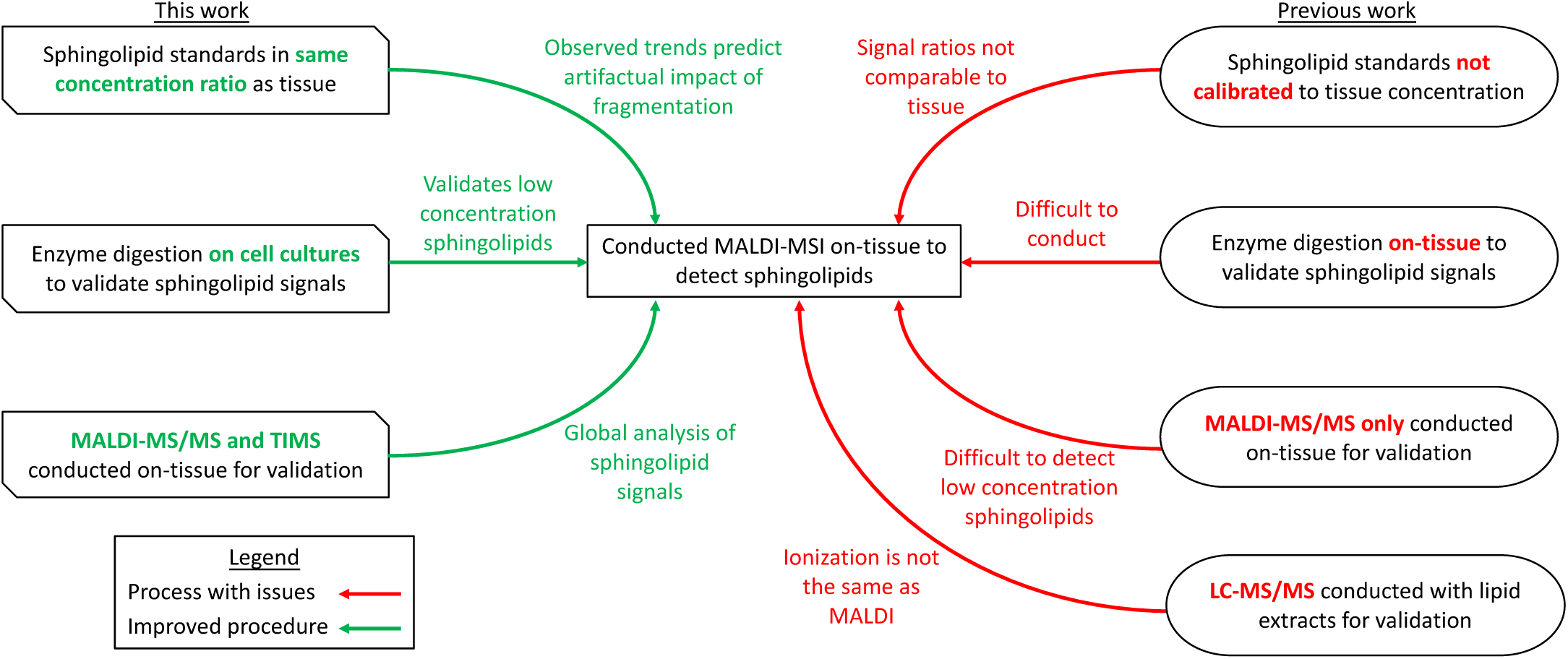
Comparison flowchart of MALDI-MSI signal detection and validation used in this study with those published in previous studies. In green, significant contributions of the presented protocol and differences with previous studies.

Furthermore, while most sphingolipids can be ionized and detected with minor/negligible fragmentation, a major source of false positive signals is the fragmentation of SM to S1P and, more profoundly C1P. In biological samples, SM levels far exceed those of S1P and C1P, making SM fragmentation a serious limitation for the measurements of these lower abundance species. Therefore, to study the impact of fragmentation of more complex and abundant sphingolipids on the simpler and less abundant species in a more meaningful way, we have utilized each standard at concentrations that reflect their relative ratios in biological samples. Additionally, we adjusted the laser power to the minimum possible level to allow the optimal trade-off between reduced fragmentation and sensitivity of detection, thereby enhancing the specificity/reliability of our signals and the contemporaneous measurement of multiple molecules. These studies thus demonstrate the severe fragmentation of SM into C1P and S1P in DAN negative mode. In the case of C1P, where the impact of SM fragmentation in the standard was massive, this event was confirmed in cells and in tissue.

1. *We modulated sphingolipids in cell culture to validate lipid signals obtained by MALDI-MSI.* In the context of MALDI, signal validation is commonly achieved by on-tissue MALDI-MS/MS. This is generally effective for high-concentration lipids, like SMs, while it can prove inadequate for low-abundance lipids, such as Cers. As summarized in **Table 1**, while a few on-tissue MALDI-MS/MS studies have focused on Cers in mouse kidney,[20, 21, 27] liver,[16] brain,[31, 34, 36] and human liver tumors [33], most MALDI-MS/MS studies focused on abundant species, particularly SMs [14, 20, 21, 23, 24, 26–30, 38–40, 55]. To address this limitation, on-tissue fragmentation of Cers has been reported utilizing the continuous accumulation of selected ions/collision-induced dissociation (CASI/CID) in the Fourier transform ion cyclotron resonance (FTICR) MALDI-MS instrument.[20] Alternatively, previous studies applied on-tissue enzymatic lipid digestion to indirectly validate the lipid signal. Application of bCDase and bSMase on kidney tissue slides from a mouse model of Farber disease was performed to confirm the identities of Cers and SMs, respectively.[20] On-tissue digestion with endoglycoceramidase validated ganglioside signals obtained with MALDI-MSI in brains from Farber mice [21, 32]. Nevertheless, enzyme application on tissue sections is not trivial to perform. We adopted a simpler approach and modulated sphingolipid levels in cell culture with enzymatic digestion (bSMase and/or pCDase) or pharmacological inhibition (eliglustat). This strategy allowed us: i) to confirm the identity of certain signals (SM, Cer, Glycosphingolipids, Sph); ii) to establish the limit of detection for certain sphingolipids as these were not detectable at basal levels (Cers, Sph, GlcCer, LacCer, C1P, S1P) and, in some cases, even after significant induction (S1P); and iii) to expand the application of MALDI-MSI to cell culture, an area with very limited studies in the literature.

Notably, using the cell culture model, we were able to validate the measurement of Sph in a biological sample, for the first time. We found the detection of intact Sph signals is achievable with minimal fragmentation of Cer, and Sph can be imaged through its water loss [M-H_2_O+H]^+^ ions at 282.279 m/z, even from tissue sections. As uncovered by the experiments using reference standards, the water-loss ion is much more abundant than the protonated [M+H]+, which is not detected in tissue. In cells, Sph was visible only after its extensive accumulation (treatment with bSMase+pCDase) where it reached a level of approximately 60 pmoles/mg of protein. This level approximates that in the PyMT (80 pmoles/mg protein) where the Sph signal is indeed registered. As far as we are aware, this is the first time a validated Sph signal is successfully reported.

Distinct signals for several GlcCer and LacCer species were present in the PyMT mammary tumor tissue sections while they were not detectable under basal conditions in the cell culture model. In this case, on-tissue validation to confirm the identity of these sphingolipids would be ideal. Our studies show that, in cells, S1P cannot be detected by MALDI-MSI neither with DHB+ mode nor with DAN-mode, even when S1P is greatly accumulated (bSMase+pCDase treatment; **Supplementary** Figure 1). On the other hand, a putative “S1P signal” is detected in DAN-mode in the PyMT tissue sample (**Supplementary** Figure 4). Since the level of S1P reached in cells after bSMase+pCDase treatment (18.8 pmoles/mg of protein) exceeds that measured in the PYMT tissue (4 pmoles/mg of protein), we are doubtful of the authenticity of the putative S1P MALDI-MSI signal registered in the PyMT. As mentioned, the signal might come from an isobaric compound (same m/z ratio).

These findings underscore the complexity and nuances involved in the MALDI imaging of sphingolipids, emphasizing the need for careful consideration of experimental conditions and methodologies. The approaches developed in this study can be utilized as a blueprint to test additional matrices and experimental conditions for further advancement of sphingolipid analysis.

## ACKNOWLEDGEMENT

This work was supported by the Stony Brook Cancer Center (SBCC) and the Lipid Signaling and Metabolism in Cancer research program. The authors wish to acknowledge the following SBCC Shared Resources (SR): the Biological Mass Spectrometry SR for expert assistance with LC/MS-MS and MALDI-MSI analysis, the Tissue Analytics SR for assisting with H&E tissue staining, and the Bioinformatics SR for assisting with signal quantitation from images and calculation of correlations. The authors are grateful to Dr. Christopher Clarke, Dr. Fabiola Velazquez, and Amira Allam for providing PyMT mammary gland tumors. The authors also thank Dr. Mehdi Damaghi and Raafat Chalar for helping with the cryosectioning of frozen tissues.

## AUTHORSHIP

B.C. performed the experiments, analyzed the data, prepared figures and wrote the manuscript; R.P. and A.G.O. designed and performed the experiments, analyzed the data, prepared tables and edited the manuscript; A.S. performed experiments and analyzed the data; S.S. analyzed the data and prepared tables and figures; X.C. assisted with bioinformatic analysis; Y.A.H. analyzed the data and edited the manuscript; C.L. and D.C. proposed the hypotheses, designed the study, analyzed the data and wrote the manuscript.

