## Supplemental Figure legends, Tables and Figures for "Critical Evaluation of Sphingolipids Detection by MALDI-MSI"

**SUPPLEMENTARY MATERIAL**

**TABLE AND FIGURE LEGENDS**

**Supplementary Table 1.** List of ion types, mass values and mass deviation of detected sphingolipid molecules**.** MALDI-MSI images were generated in Python with a mass deviation of 10ppm from the mass value found on LIPIDMAPS corresponding to each respective sphingolipid species and ion type.

**Supplementary Table 2.** MALDI signal of different sphingolipid standards, normalized by internal standards, mixed with DAN or DHB matrices and evaluated in negative (-) or positive (+) ion detection mode. ND: Not determined.

**Supplementary Table 3.** Assessing fragmentation of authentic standards in DHB and DHA positive ion detection mode. Ratios between the signal of the produced fragments and the relative abundance of the same fragment molecules in biological samples were derived using values from the fragment signal column reported in this Table and signal values in Table 2. Values in Table 2 and this Supplementary Table were collected in the same experiments. DHB signals were acquired in positive ion detection mode and 35% laser power. DHA signals were acquired in positive ion detection mode and 30% laser power. Color scale red to yellow indicates high to low fragment production (ratio <0.1 indicates that production of fragment has a Minor impact; ratio 0.1-0.2 indicates that production of fragment has a Mild impact; ratio >0.2 and <0.6 indicates that production of fragment has a Significant impact; ratio >0.6 indicates that production of fragment has a Severe impact).

**Supplementary Figure 1.** LC-MS/MS sphingolipid measurements in parallel to MALDI-MSI analysis upon bSMase and pCDase treatments. HeLa cells were grown and treated in 60mm dishes. Immediately after treatment, cells were quenched with lipid extraction mixture and lipid extraction and LC-MS/MS analysis were performed as indicated in Materials and Methods. Statistics: One-way ANOVA with post hoc Dunnett's multiple comparisons test compared to CT. ∗P<0.01; ∗∗P<0.001; ∗∗∗P<0.001; ∗∗∗∗P<0.0001.

**Supplementary Figure 2.** MALDI-MSI images of major SMs, detected in positive ion mode with DHB matrix on a section of PyMT mammary tumor.  Shown are the protonated, sodium, and potassium adduct ions, as well as combined positive ions are.

**Supplementary Figure 3.** MS/MS on-tissue fragmentation of [SM(d18:1/16:0)+H]+ (red) in comparison to MS/MS fragmentation of C16-SM standard (blue) on the mass-to-charge ratio range of (**A**) 50-750 m/z and (**B**) 200-750 m/z. Characteristic ions at 184.073 m/z and 264.267 m/z were found for SMs.

**Supplementary Figure 4**. MALDI-MSI images of S1P, C1P and C16-SM deprotonated species detected in negative ion mode with DAN matrix, along with C16-SM protonated specie detected in positive ion mode with DHB matrix from consecutive sections of PyMT mammary tumor.

**Supplementary Figure 5.** Pearson correlation cluster map of sphingolipid species from PyMT tissue sample. S1P is highlighted in red showing no strong correlation with SM.

| Sphingolipid Species | Ion Type | Mass Value (m/z) | Mass Deviation (m/z) |
|---|---|---|---|
| **Sph** | $[M+H-H_2O]^+$ | 282.279 | 0.003 |
| **Sph** | $[M+H]^+$ | 300.290 | 0.003 |
| **C16:0-SM** | $[M+H]^+$ | 703.575 | 0.007 |
| **C24:1-SM** | $[M+H]^+$ | 813.684 | 0.008 |
| **C24:0-SM** | $[M+H]^+$ | 815.700 | 0.008 |
| **S1P** | $[M-H]^-$ | 378.242 | 0.004 |
| **C16:0-C1P** | $[M-H]^-$ | 616.471 | 0.006 |
| **C16:0-Cer** | $[M-H]^-$ | 536.505 | 0.005 |
| **C24:1-Cer** | $[M-H]^-$ | 646.614 | 0.006 |
| **C24:0-Cer** | $[M-H]^-$ | 648.630 | 0.006 |
| **C16:0-GlcCer** | $[M-H]^-$ | 698.558 | 0.007 |
| **C24:1-GlcCer** | $[M-H]^-$ | 808.667 | 0.008 |
| **C24:0-GlcCer** | $[M-H]^-$ | 810.683 | 0.008 |
| **C16:0-LacCer** | $[M-H]^-$ | 860.611 | 0.009 |
| **C24:1-LacCer** | $[M-H]^-$ | 970.720 | 0.010 |
| **C24:0-LacCer** | $[M-H]^-$ | 972.736 | 0.010 |
| **C16:0-GM3** | $[M-H]^-$ | 1151.706 | 0.012 |
| **C18:0-GM3** | $[M-H]^-$ | 1179.737 | 0.012 |
| **C24:1-GM3** | $[M-H]^-$ | 1261.815 | 0.013 |
| **C24:0-GM3** | $[M-H]^-$ | 1263.831 | 0.013 |
| **C16:0-GM1** | $[M-H]^-$ | 1516.838 | 0.015 |
| **C18:0-GM1** | $[M-H]^-$ | 1544.869 | 0.015 |
| **C24:1-GM1** | $[M-H]^-$ | 1626.948 | 0.016 |
| **C24:0-GM1** | $[M-H]^-$ | 1628.963 | 0.016 |

**Supplementary Table 1**. List of ion types, mass values and mass deviation of detected sphingolipid molecules.

| Sphingolipid Standards | Concentrations tested (uM) | Replicate #1 | | Replicate #2 | | Replicate #3 | |
|---|---|---|---|---|---|---|---|
| | | DAN (-) | DHB (+) | DAN (-) | DHB (+) | DAN (-) | DHB (+) |
| C16-SM | 5000 | 5.79 | 27.03 | 9.18 | 50.39 | 5.38 | 26.65 |
| C16-Cer | 50 | 1.35 | 0.60 | 1.52 | 0.37 | 1.61 | 0.50 |
| C16-GlcCer | 50 | 0.65 | 0.14 | 0.32 | 0.13 | 0.97 | 0.17 |
| C16-LacCer | 50 | 0.756 | 0.057 | 0.501 | 0.033 | 1.74 | 0.018 |
| C18-Cer | 50 | 1.27 | 0.193 | 1.32 | 0.130 | 1.26 | 0.339 |
| C18-GlcCer | 50 | 1.74 | 0.304 | 1.51 | 0.191 | 1.42 | 0.157 |
| C18-LacCer | 50 | 2.580 | 0.0282 | 0.474 | 0.0528 | 0.376 | 0.0345 |
| GM1 | 50 | 1.37 | 1.07 | 1.75 | 1.22 | 1.37 | 0.874 |
| GM3 | 50 | 2.54 | 0.787 | 4.30 | 1.14 | 2.45 | 1.12 |
| Sphingosine | 10 | 0.267 | ND | 0.268 | ND | 0.162 | ND |
| C16-C1P | 1 | 0.060 | 0.119 | 0.016 | 0.100 | 0.053 | 0.126 |
| S1P | 1 | 0.468 | 0.220 | 0.419 | 0.137 | 0.424 | 0.252 |

**Supplementary Table 2**. MALDI signal of different sphingolipid standards, normalized by internal standards, mixed with DAN or DHB matrices and evaluated in negative (-) or positive (+) ion detection mode.

| DAN negative ion detection mode | Replicate #1 | | | Replicate #2 | | | Replicate #3 | | |
|---|---|---|---|---|---|---|---|---|---|
| **Potential fragmentation path: parent to fragment** | Fragment signal | Fragment signal/ Relative physiologic levels of fragment molecule | Impact of fragment on relative physiologic levels | Fragment signal | Fragment signal/ Relative physiologic levels of fragment molecule | Impact of fragment on relative physiologic levels | Fragment signal | Fragment signal/ Relative physiologic levels of fragment molecule | Impact of fragment on relative physiologic levels |
| C16-SM → S1P | 1357 | 1.9 | Severe | 760 | 5 | Severe | 664 | 4.4 | Severe |
| C16-S1P → Sphingosine | 293 | 0.36 | Significant | 73 | 0.12 | Mild | 149 | 0.36 | Significant |
| C16-C1P → C16-Cer | 92 | 0.0010 | Negligible | 114 | 0.0048 | Negligible | 87 | 0.0021 | Negligible |
| **DHB positive ion detection mode** | Replicate #1 | | | Replicate #2 | | | Replicate #3 | | |
| **Potential fragmentation path: parent to fragment** | Fragment signal | Fragment signal/ Relative physiologic levels of fragment molecule | Impact of fragment on relative physiologic levels | Fragment signal | Fragment signal/ Relative physiologic levels of fragment molecule | Impact of fragment on relative physiologic levels | Fragment signal | Fragment signal/ Relative physiologic levels of fragment molecule | Impact of fragment on relative physiologic levels |
| C16-SM → C16-Cer | 12 | 0.0028 | Negligible | 7 | 0.0011 | Negligible | 3 | 0.002 | Negligible |
| C16-GlcCer → C16-Cer | 223 | 0.052 | Minor | 483 | 0.079 | Minor | 793 | 0.52 | Significant |
| C16-LacCer → C16-Cer | 591 | 0.14 | Mild | 870 | 0.14 | Mild | 823 | 0.54 | Significant |
| C16-C1P → C16-Cer | 17 | 0.0047 | Negligible | 8 | 0.0013 | Negligible | 11 | 0.0075 | Negligible |
| C16-S1P → Sphingosine | 1790 | 0.17 | Mild | 661 | 0.064 | Minor | 405 | 0.044 | Negligible |
| C18-GM1 → C18-Cer | 235 | 0.064 | Minor | 3 | 0.00044 | Negligible | 10 | 0.0066 | Negligible |
| C18-GM1 → C18-GlcCer | 807 | 0.22 | Significant | 238 | 0.23 | Significant | 225 | 0.35 | Significant |
| C18-GM1 → C18-LacCer | 280 | 2.0 | Severe | 113 | 1.0 | Severe | 116 | 0.73 | Severe |
| C18-GM1 → C18-GM3 | 783 | 0.32 | Significant | 232 | 0.85 | Severe | 202 | 0.33 | Significant |
| C18-GM3 → C18-Cer | 286 | 0.078 | Minor | 23 | 0.0038 | Negligible | 15 | 0.0099 | Negligible |
| C18-GM3 → C18-GlcCer | 2790 | 1.7 | Severe | 300 | 0.29 | Significant | 1145 | 1.8 | Severe |
| C18-GM3 → C18-LacCer | 1146 | 8.1 | Severe | 191 | 1.8 | Severe | 313 | 2.0 | Severe |
| **DHA positive ion detection mode** | Replicate #1 | | | Replicate #2 | | | Replicate #3 | | |
| **Potential fragmentation path: parent to fragment** | Fragment signal | Fragment signal/ Relative physiologic levels of fragment molecule | Impact of fragment on relative physiologic levels | Fragment signal | Fragment signal/ Relative physiologic levels of fragment molecule | Impact of fragment on relative physiologic levels | Fragment signal | Fragment signal/ Relative physiologic levels of fragment molecule | Impact of fragment on relative physiologic levels |
| C16-SM → C16-Cer | 21 | 0.0017 | Negligible | 23 | 0.0016 | Negligible | 4 | 0.00071 | Negligible |
| C16-GlcCer → C16-Cer | 2677 | 0.22 | Significant | 5099 | 0.37 | Significant | 1963 | 0.38 | Significant |
| C16-LacCer → C16-Cer | 8371 | 0.68 | Severe | 11335 | 0.82 | Severe | 4353 | 0.84 | Severe |
| C16-C1P → C16-Cer | 119 | 0.0096 | Minor | 88 | 0.0064 | Negligible | 33 | 0.0063 | Negligible |
| C16-S1P → Sphingosine | 3263 | 4.2 | Severe | 1819 | 0.017 | Minor | 2013 | 0.038 | Minor |
| C18-GM1 → C18-Cer | 66 | 0.0053 | Negligible | 8.7 | 0.00063 | Negligible | 26 | 0.0051 | Negligible |
| C18-GM1 → C18-GlcCer | 1579 | 0.39 | Significant | 208 | 0.24 | Significant | 989 | 0.53 | Significant |
| C18-GM1 → C18-LacCer | 366 | 0.43 | Significant | 142 | 0.32 | Significant | 381 | 0.68 | Significant |
| C18-GM1 → C18-GM3 | 1789 | 0.15 | Mild | 318 | 0.23 | Significant | 1213 | 0.21 | Significant |
| C18-GM3 → C18-Cer | 106 | 0.0085 | Negligible | 21 | 0.0015 | Negligible | 31 | 0.0059 | Negligible |
| C18-GM3 → C18-GlcCer | 8308 | 2.1 | Severe | 1194 | 1.4 | Severe | 4326 | 2.3 | Severe |
| C18-GM3 → C18-LacCer | 3515 | 4.1 | Severe | 554 | 1.2 | Severe | 1801 | 3.2 | Severe |

**Supplementary Table 3.** Assessing fragmentation of authentic standards in DHB and DHA positive ion detection mode.

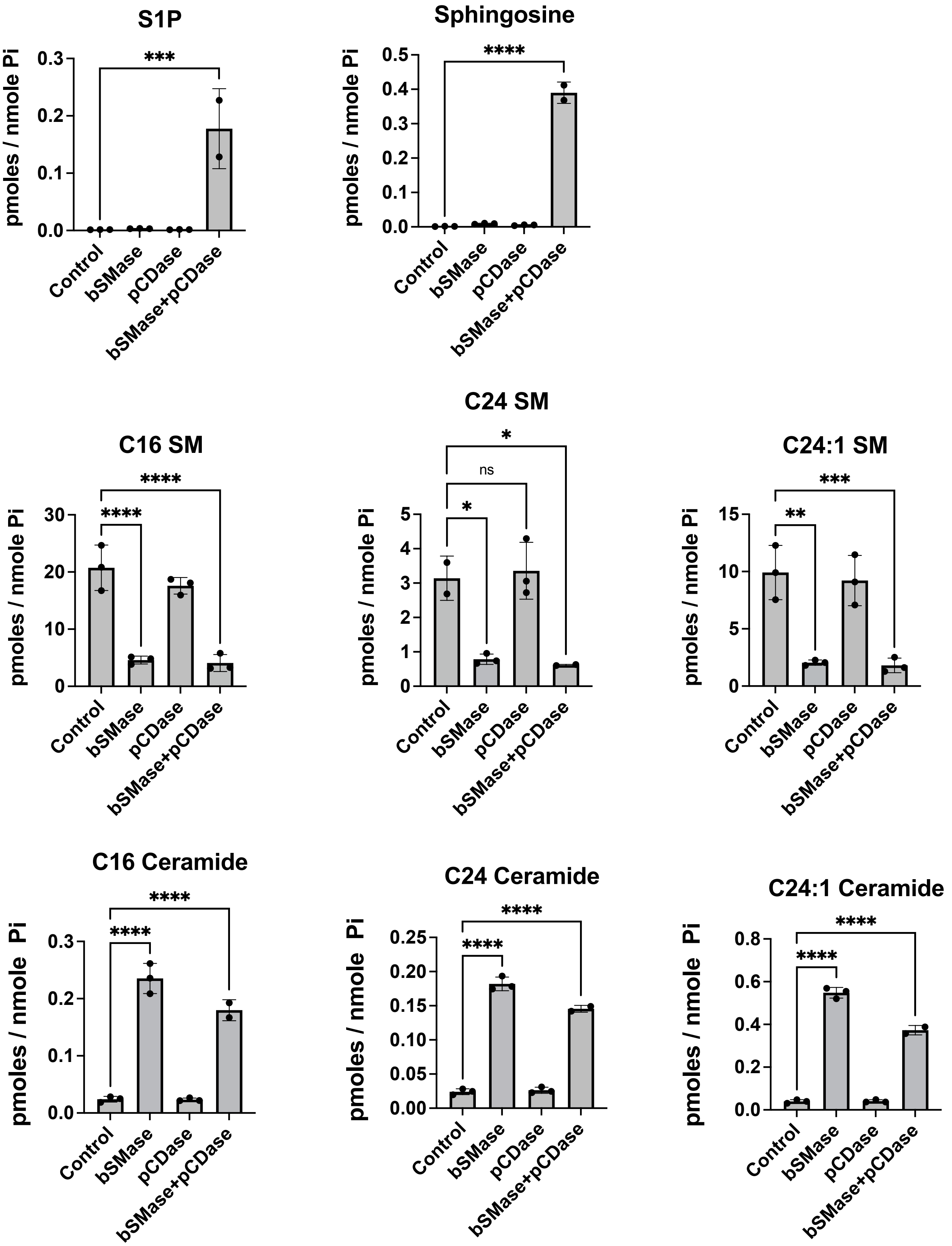


**Supplementary Figure 1**.

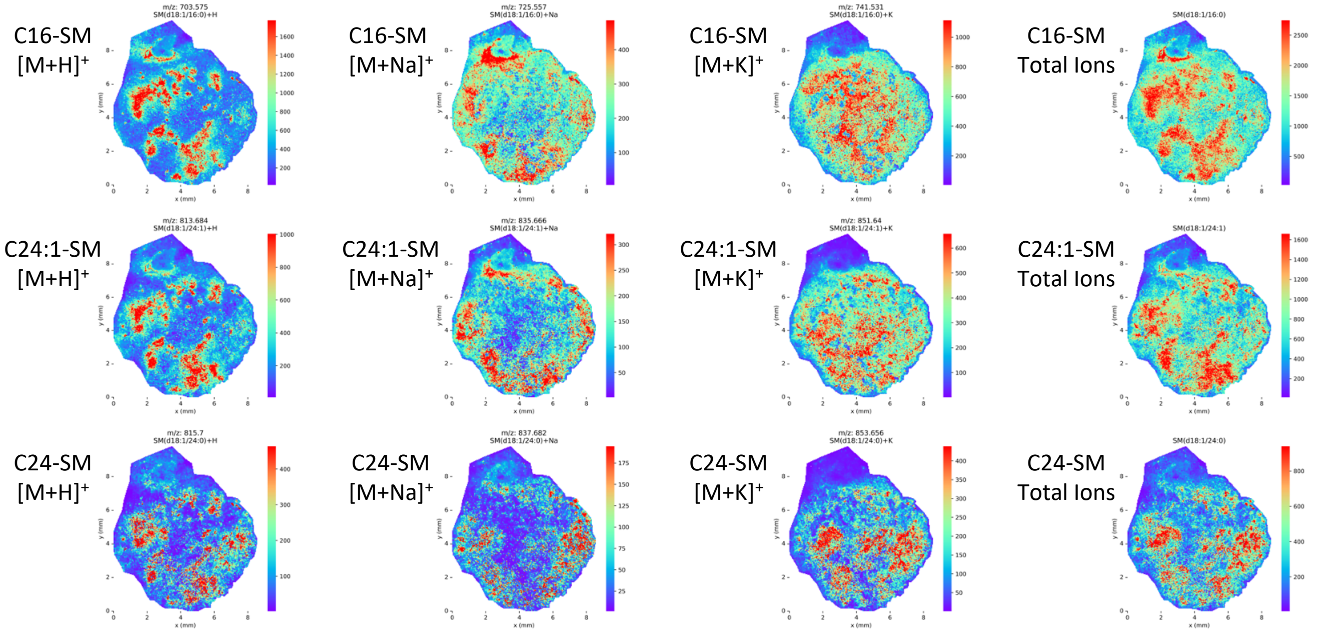


**Supplementary Figure 2**.

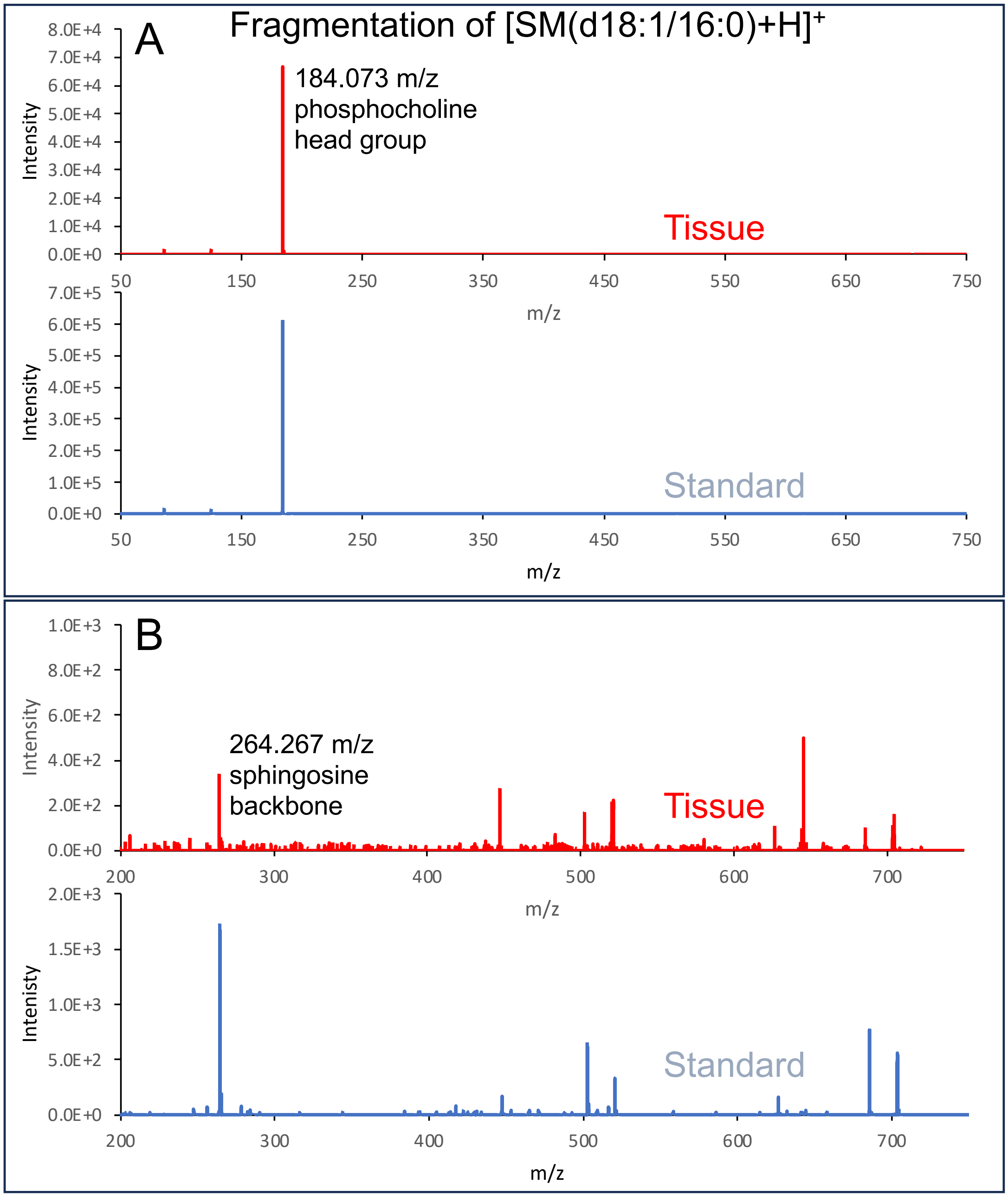


**Supplementary Figure 3**.

378.242 m/z
S1P [M-H]$^-$ ?

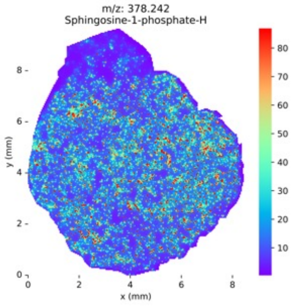


616.471 m/z
C16-C1P [M-H]$^-$ ?

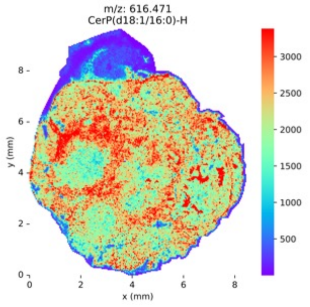


701.56 m/z
C16-SM [M-H]$^-$ ?

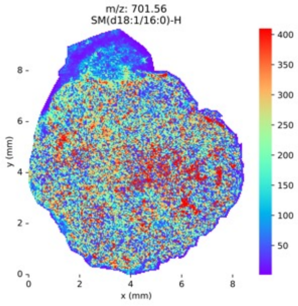


703.575 m/z
C16-SM [M+H]$^+$

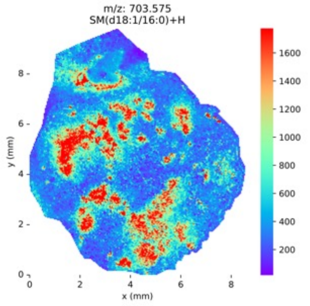


**Supplementary Figure 4**.

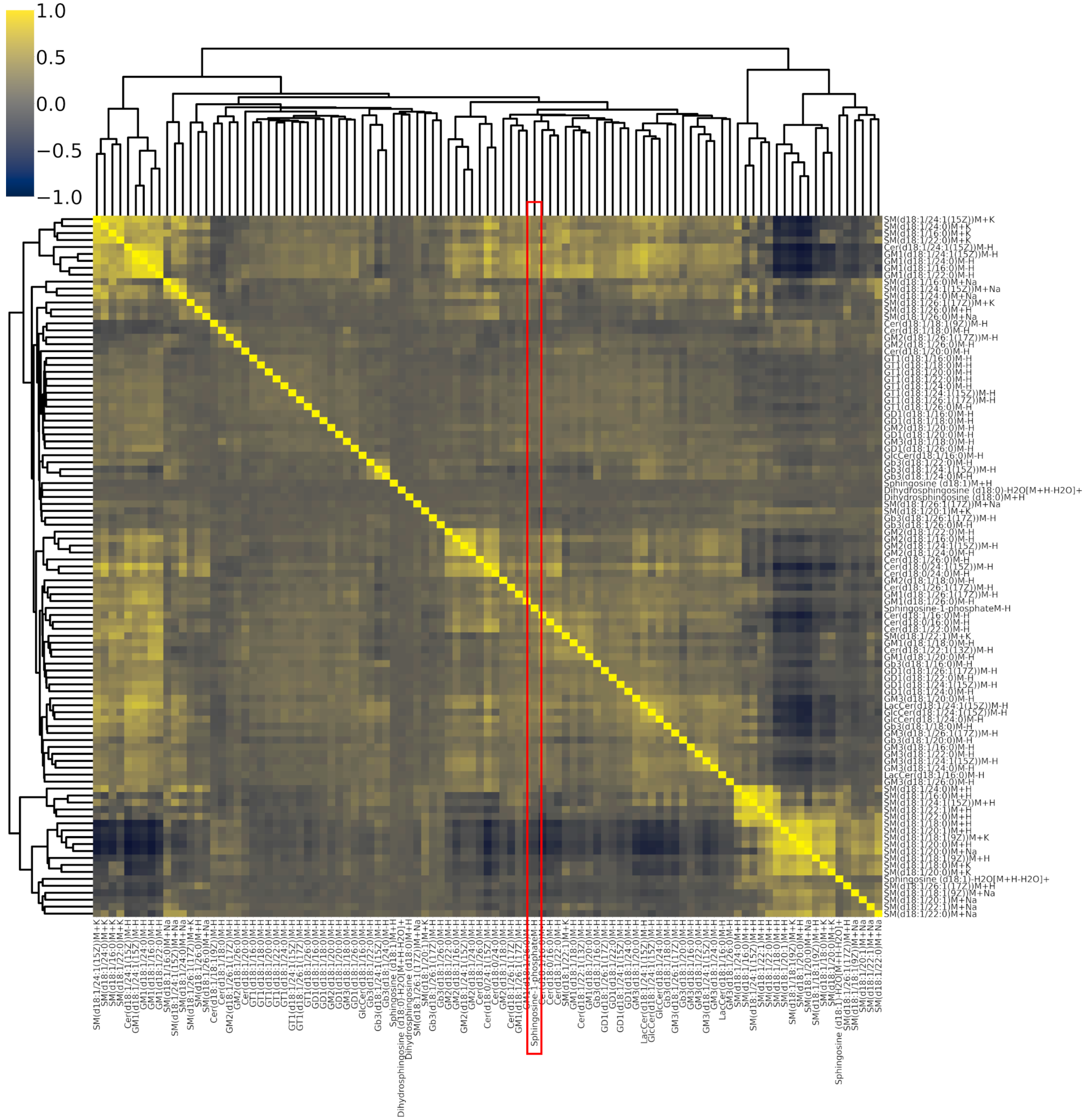


**Supplementary Figure 5**.
